# Dual Drug Targeting to Kill Colon Cancer Cells

**DOI:** 10.1101/2021.02.28.433288

**Authors:** Silvia Paola Corona, Francesca Walker, Janet Weinstock, Guillaume Lessene, Maree Faux, Antony W Burgess

**Affiliations:** Structural Biology Division, WEHI, 1G Royal Parade, Parkville, Australia, 3052; Personalised Oncology Division, WEHI, 1G Royal Parade, Parkville, Australia, 3052; Department of Medical Biology, University of Melbourne, Parkville, Australia, 3052; Department of Surgery, Royal Melbourne Hospital, The University of Melbourne, Parkville, Australia 3052; Ludwig Institute for Cancer Research, Parkville, Australia, 3052; Chemical Biology Division, WEHI, 1G Royal Parade, Parkville, Australia, 3052; Department of Pharmacology and Therapeutics, The University of Melbourne, Parkville, Australia 3052

## Abstract

Colorectal cancer (CRC) is driven by a small set of oncogenic and tumor suppressor mutations. However, different combinations of mutations often lead to poor tumor responses to individual anti-cancer drugs. We have investigated the anti-proliferative and *in vitro* cytotoxic activity of pair-wise combinations of inhibitors which target specific signalling pathways. Colon cancer cells in non-adherent cultures were killed more effectively by combinations of pyrvinium pamoate (a Wnt pathway inhibitor) and ABT263 (a pro-apoptotic Bcl-2 family inhibitor) or Ly29004 (a PI3kinase inhibitor). However, in a mouse xenograft model, the formulation and toxicity of the ABT737/PP combination prevent the use of these drugs for treatment of tumors. Fortunately, oral analogues of PP (pyrvinium phosphate, PPh) and ABT737(ABT263) have equivalent activity and can be used for treatment of mice carrying SW620 colorectal cancer xenografts. The PPh/ABT263 induced SW620 tumor cell apoptosis and reduced the rate of SW620 tumor growth. Combinations of Wnt signaling inhibitors and specific inhibitor of pro-survival proteins should be considered for the treatment of precancerous colon adenomas and advanced colorectal cancers with APC mutations.

## Introduction

The treatment of colorectal cancer has improved considerably over the last decade^1^, however, while the use of targeted therapies, e.g. epidermal growth factor receptor inhibitors and antibodies^2^, has improved the progression free survival for patients with advanced colorectal cancer (CRC) ^3^, the 5-years survival rates have only improved marginally. The genetics of CRC have been analysed in considerable detail and a small number of oncogenic mutations (APC, TP53, Kr-RAS, PI3K, RNF-43, RSPO fusions, B-RAF, β-catenin, TGF β receptor, SMAD 2,3,4 and the mismatch repair enzymes) drive this cancer ^4^, thus there is sufficient variation caused by different combinations and locations of these mutations to make CRC a genetically diverse disease. Targeting a single mutation may slow the proliferation of CRC cells and in some cases may even kill the cells, however, in most cases the CRC cells will escape targeted therapies such as cetuximab ^5^.

We have selected a small panel of CRC cell lines with a range of oncogenic and tumor suppressor mutations and measured the sensitivity of these cells to agents which inhibit a single oncogenic target or signalling pathway associated with CRC. We have used assays which measure the ability of the drugs to inhibit proliferation and/or kill these CRC cell lines *in vitro*. Once we determined the sensitivity of the cell lines to single inhibitors, we tested the ability of pairwise combinations of inhibitors to kill specific CRC genotypes. Depending on the mutational profile we identified dual targeting drug combinations which killed CRC cells at low concentrations *in vitro*. The most effective combination involved the inhibition of Wnt signalling^6^ and the induction of apoptosis by inhibiting the pro-survival protein Bcl-2^7^. Given that most CRC are initiated by the loss of APC-function a Wnt^i^/Bcl-2^i^ drug combination was also explored in a mouse xenograft model of colon cancer using SW620 cells. The use of drug combinations to target the signaling pathways perturbed during the initiation of progression of CRC has potential for both preventing CRC and improving outcomes for patients^8^.

## Materials and Methods

### Antibodies and Reagents

Antibodies were obtained from BD Transduction Laboratories (mouse monoclonal anti-β-catenin and mouse monoclonal anti-E-Cadherin), Cell Signaling Technology (rabbit polyclonal anti-S33/S37/T41 phospho β-catenin), Sigma-Aldrich (mouse monoclonal anti-β-tubulin), Abcam (rabbit polyclonal anti-Lamin B1), Li-COR Biosciences (IRDye 800CW Goat anti-rabbit; IRDye 800CW Goat anti-mouse). Propidium Iodide, Hoechst 33342 for live cell imaging and MTT were purchased from Sigma Aldrich (St. Louis). The LDH Cytotoxicity Detection Kit was purchased from Roche Diagnostics, Mannheim. Pyrvinium Pamoate was purchased from USP, Rockville, MD, USA. Pyrvinium Phosphate was synthesized at the WEHI, Bundoora, VIC, following the protocol published by Yu and colleagues^9^. ABT737 was purchased from SYNTHESIS MED CHEM, South Yarra, VIC, Australia. ABT263 (Navitoclax) was purchased from CAPOT Chemicals, Shanghai. DAPT was purchased from TOCRIS Bioscience, Ellisville. The EGFR small molecule inhibitor AG1478 mesylate was purchased by from the Institute of Drug Technology (IDT, Boronia). The PI3K inhibitor LY294002 was purchased from Calbiochem (Merck Chemicals, Darmstadt); the SRC-kinase family inhibitor (WEHI-1208800) was synthesized at WEHI, Bundoora according to the Patent Application No. PCT/AU2011/000858.

The ApopTag Peroxidase *In Situ* Apoptosis Detection Kit was purchased from Millipore (Billerica)

### Cell culture

SW620 were a kind gift of Prof. John Mariadason, Ludwig Institute for Cancer Research, Heidelberg, VIC. LIM1899 and LIM2537 colorectal cancer cell lines are available WEHI, Parkville. The cell lines were maintained in RPMI-1640 medium (GIBCO) supplemented with 10% v/v Fetal Bovine Serum (GIBCO), Thioglycerol (10μM final concentration), Insulin (2.5 U/100ml), Hydrocortisone (0.1mg/100ml) and antibiotics (Penicillin 0.6g/100ml and Streptomycin 1g/100ml), at 37°C and 5% CO2 in humidified atmosphere. Cells were validated by DNA sequencing^4^.

### Drug Formulations

The stock solutions of the drugs were dissolved in the vehicles suggested by previous reports^10, 11, 12, 9^. Briefly, Pyrvinium Pamoate (PP) was dissolved in 2% DMSO-saline(v/v) (final concentration 0.5 mg/ml); Pyrvinium Phosphate was dissolved in water (final concentration 0.5 mg/ml); ABT737 was dissolved in 30% v/v Propylene Glycol, 65% v/v D5W (5%w/v dextrose in distilled water), 5% v/v Tween 80 (final concentration 5 mg/ml); ABT263 was dissolved in 30% v/v Polyethylene Glycol, 60% v/v Phosal 50 PG (50% w/v phosphatidylcholine in propylene glycol used to enhance solubility and bioavailability of compounds), 10% v/v Ethanol (final concentration 5mg/ml).

### Mice

-6 to 8 weeks Non-obese Diabetic with Severe Combined Immunodeficiency NOD/SCID (NODCB17-*Prkdcscid*/ARC) mice were obtained from the Animal Resources Centre (ARC, Perth). All animal procedures were approved and carried out in accordance with the Animal Ethics Committees of the Ludwig Institute for Cancer Research, Melbourne Branch or WEHI.

### MTT Assay

Unless otherwise specified, cells were plated at 10^4^ cells/well in 100μl of medium (RPMI-1640 plus Fetal Bovine Serum 5%) for the assay and incubated overnight at 37°C in 5% CO2 in a humidified atmosphere. 150μl/well of RPMI 1640 with Fetal Bovine Serum (FBS) 5% was aliquoted into each well of the 96well plate and 150μl/well of the 4x concentration inhibitor was added to the first well of each row to obtain a concentration 2x the one set as starting concentration of the experiment. A serial 2-fold dilution was then performed across the plate. After 3-4 days of incubation, MTT (Sigma-Aldrich, St. Louis) 10μl/well was added to the plates and cells were incubated for 4 hours at 37°C following the manufacturer’s instructions. Plates were then centrifuged at 1500rpm for 10 minutes to collect all the cells at the bottom of the wells. Medium was removed carefully and acidified isopropanol (0.04M HCl in Isopropanol) was added at 200μl/well to solubilize the purple formazan crystals. Plates were shaken on a Vibramax 100 plate shaker (Heidolph Instruments, Kelheim) for 30 minutes at 450rpm to speed the solubilization process. The Optical Density at 560/690nm was measured on a Multiskan Ex Spectrophotometer (Thermofisher Scientific, Waltham).

### The LDH Cytotoxicity Detection Assay

(Roche Diagnostics GmbH, Roche Applied Science, Mannheim, Germany). To assess the doubling time of each cell line and the optimal number of cells to plate for the experiments, under both adherent and anchorage independent conditions (“hanging drops”), a cell titration experiment was performed for each cell line on cells grown under adherent conditions or organoids formed in hanging drops.

### LDH Assay - adherent conditions

Cells were harvested at 80% confluence after incubation for 5 minutes in a solution containing 0.1% w/v Trypsin in Versene 0.02% w/v. After centrifugation at 1500rpm for 5 minutes, cells were re-suspended in fresh medium and viability and cell numbers were monitored with vital dye Trypan Blue 0.2% w/v (Trypan Blue 0.4% w/v, Sigma-Aldrich, St. Louis). The starting number of cells for the titration experiments on cells cultured under adherent conditions was set at 2 × 10^4^ cells in 100μl of culture medium. 100μl of 5% Fetal Bovine Serum (FBS) in RPMI 1640 was aliquoted into each well of a 96 well plate. 100μl of cells at 2x the set starting number were added to the first rows of the plate and a 2-fold serial dilution was performed across the plate. At the end of the titration, the volume in each well was 100μl, with 6 wells of each cell dilution. The last two rows of the 96well plate were used for the medium alone (background), to be used for the normalization of the results. For inhibitor studies, cells were plated on 96well-plates at 5000 cells/well, the plates were incubated overnight at 37°C, 5% CO2 in humidified atmosphere. After 24 hours, drugs were diluted to the starting concentration and the respective dilutions of the drugs were then added to the cells across the plate to reach the desired concentration, and plates were incubated for 72 hours and processed as per the manufacturer’s instructions.

### LDH Assay - anchorage independent conditions

Cells were grown under anchorage independent conditions using the “hanging drops” method (Robinson et al 2004) adapted to 96-well plates. The starting cell number was set at 10^5^ cells per drop (30μl/drop in RPMI-1640 with Fetal Bovine Serum, FBS, 5%). A 2-fold serial dilution of drugs was then added to the cells.

### Xenografts

NOD/SCID female and male mice, 9 to 10 weeks of age, were used for the xenograft experiments. The mice were divided randomly in groups, 8 mice/experimental group, and the mice from each experimental group were placed in two cages, 4 mice per cage. SW620 colorectal cancer cells were harvested by trypsinization, washed in serum free medium and resuspended in PBS at 5 x10^7^/ml. SW620 cells were inoculated in 100μl of PBS at 5 x10^6^ cells/tumor, 2 tumors per mouse, subcutaneously, on the left and right flanks. Mice were inoculated under general anesthesia with isofluorane. Administration of the drugs was started on the 8th day after cell inoculation. The solutions were administered by gavage (final volume 200μl). For combination therapy the 2 drug administrations per day were given with at least a 3 hour interval between each other. For the controls the vehicles were administered without the compounds. Mice were weighed and checked for health problems, and tumor volume measured by calliper in two dimensions, at least twice a week, with the mice under a light general anesthesia to increase the accuracy of the measurements. The mice were euthanized after 3 to 4 weeks of treatment or when the tumor volume had reached ethically unacceptable size. All the tumors were harvested and weighed at the end of the experiment and preserved in 10% v/v buffered formalin for histological analysis. 4 mice/group were randomly chosen and the spleens, livers and kidneys from 4 mice/group chosen randomly were preserved in buffered 10% v/v formalin for histological analysis.

### Histology

Detection of apoptosis *in vivo* was performed using the ApopTag Peroxidase *In Situ* Apoptosis Detection Kit (Millipore, Billerica, MA, USA) according to the manufacturer’s instructions.

### Analysis and quantitation of apoptosis

Images of the slides obtained with the ApopTag assay were acquired digitally with the Aperio ScanScope XT (Vista). 20 fields/image/tumor were chosen arbitrarily within the epithelial tumor tissue, away from the stromal component and from the margins of the sections. Brown-stained cells were manually counted within each field. We used MetaMorph version 7.7.10 (Molecular Devices, Sunnyvale) to count the total and dead cells fields for each tumor and to calculate the area of positively stained cells as well as the total area. The results were then expressed as apoptotic cells area over the total area.

### Statistics

Statistical analysis were performed using GraphPad Software-Unpaired *t*-test. Graphs were plotted using GraphPad Prism v.6, Origin or Matplotlib. All *in vitro* experiments were done in triplicate and each data point was calculated from triplicate wells.

## Results

### Single agent inhibition of proliferation

In order to measure proliferation of cells after the exposure to small molecule pathway inhibitors and to quantify the relative dependency of the cells on each signaling pathway, three colon cancer cell lines, SW620, LIM1899 and LIM2537 (see cell line genetic characteristics in Supplementary Table 1) were treated with increasing concentrations of signaling inhibitors: AG1478, WEHI-1208800, LY294002, ABT737, PP or DAPT (see Supplementary Table 2 for details of these inhibitors). The results of these experiments are summarised in Table 1 and Supplementary Figure 1; we note that DAPT had no effect on the proliferation or killing of any of these cell lines even up to 10μM, so the results were not included in the table. The EGFR inhibitor AG1478 totally abolished proliferation of LIM1899 cells, but, even at the highest concentrations (> 10μM) it showed only a partial cytostatic effect on LIM2537 (Table 1). SW620 cell line is also resistant to EGFR inhibition by AG1478, however, this is expected, as these cells do not express the EGF receptor ^13,14^. Clearly, the LIM1899 cell line is dependent on the EGFR signaling pathway for proliferation *in vitro* (Table 1); this result is rather unexpected given that these cells harbor a KRAS mutation, which, in the majority of colorectal tumors, circumvents the EGF signaling pathway inhibition in the clinic ^15^.

Pyrvinium Pamoate (PP) is a recognized inhibitor of Wnt signaling ^16-18^. We confirmed that PP (2µM) inhibited Wnt signaling using a TCF-driven-GFP reporter system in both LIM1899 and SW620 cells (Supplementary Figure 2). Although PP inhibits Wnt signaling, it has been reported to inhibit other targets ^9^. There was no correlation between the genotype of the cell lines tested and their sensitivity to PP: the drug potently inhibited the proliferation of all 3 cell lines at low concentrations (50% cytostatic effect at around 50nM, Table 1 and Supplementary Figure 1), suggesting that Wnt signaling pathway activation is required for proliferation of these cells *in vitro*, independently of whether they carry an APC or a β-catenin mutation.

**Table 1:**
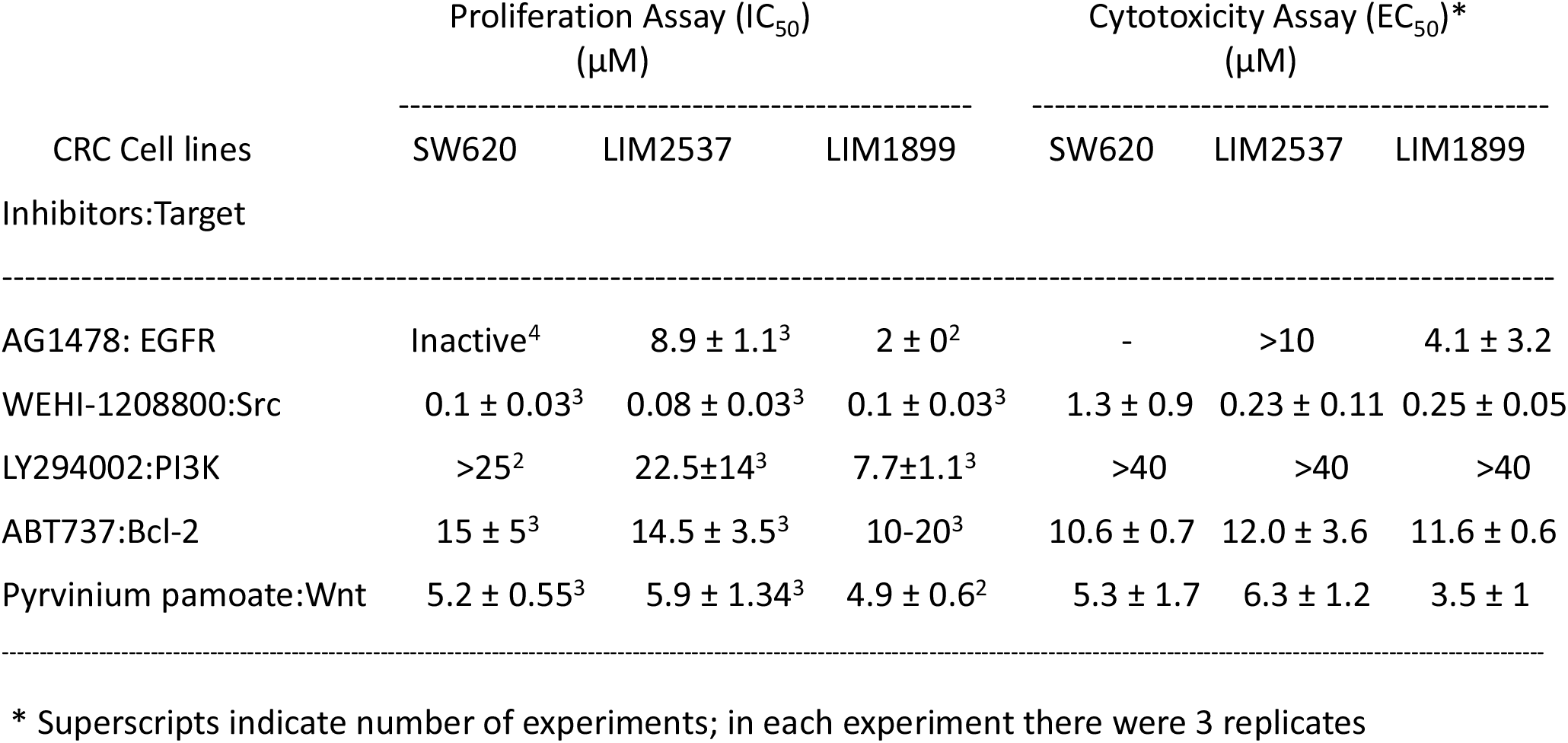
Effects of signaling inhibitors on proliferation and viability of three colorectal cancer cell lines in adherent cultures

The potent inhibition of LIM1899 cells by Pyrvinium Pamoate (PP) was unexpected: this cell line is characterized by the presence of a point mutation in β-catenin which constitutively activates the Wnt pathway and renders these cells resistant to the extracellular Wnt-inhibitor Dickkopf-1 (DKK-1) ^19^. If PP exerts its only inhibitory function through CK-1α, the mutation which alters β-catenin at the Ser45 residue should render the cells resistant to inhibition of the Wnt pathway by PP. The inhibition of LIM1899 by Pyrvinium Pamoate strongly suggests that the inhibitory activity of the compound is either exerted on components of the Wnt signaling pathway downstream of β-catenin, or that the drug acts through GSK3 activation, as suggested by Venerando et al. ^16^ or that the drug affects cell viability independently of the Wnt pathway.

The SRC-Family kinases Inhibitor WEHI-1208800 also inhibits the proliferation of all three cell lines within a similar IC_50_ range (100nM) (Table 1, Supplementary, Figure 1). This data suggests a strong degree of dependency of these cells on active SRC signaling pathway for proliferation *in vitro*, however, as for most kinase inhibitors, cross reactivity mean that this specificity is only a potential indication of its mechanism of action.

**Figure 1:**
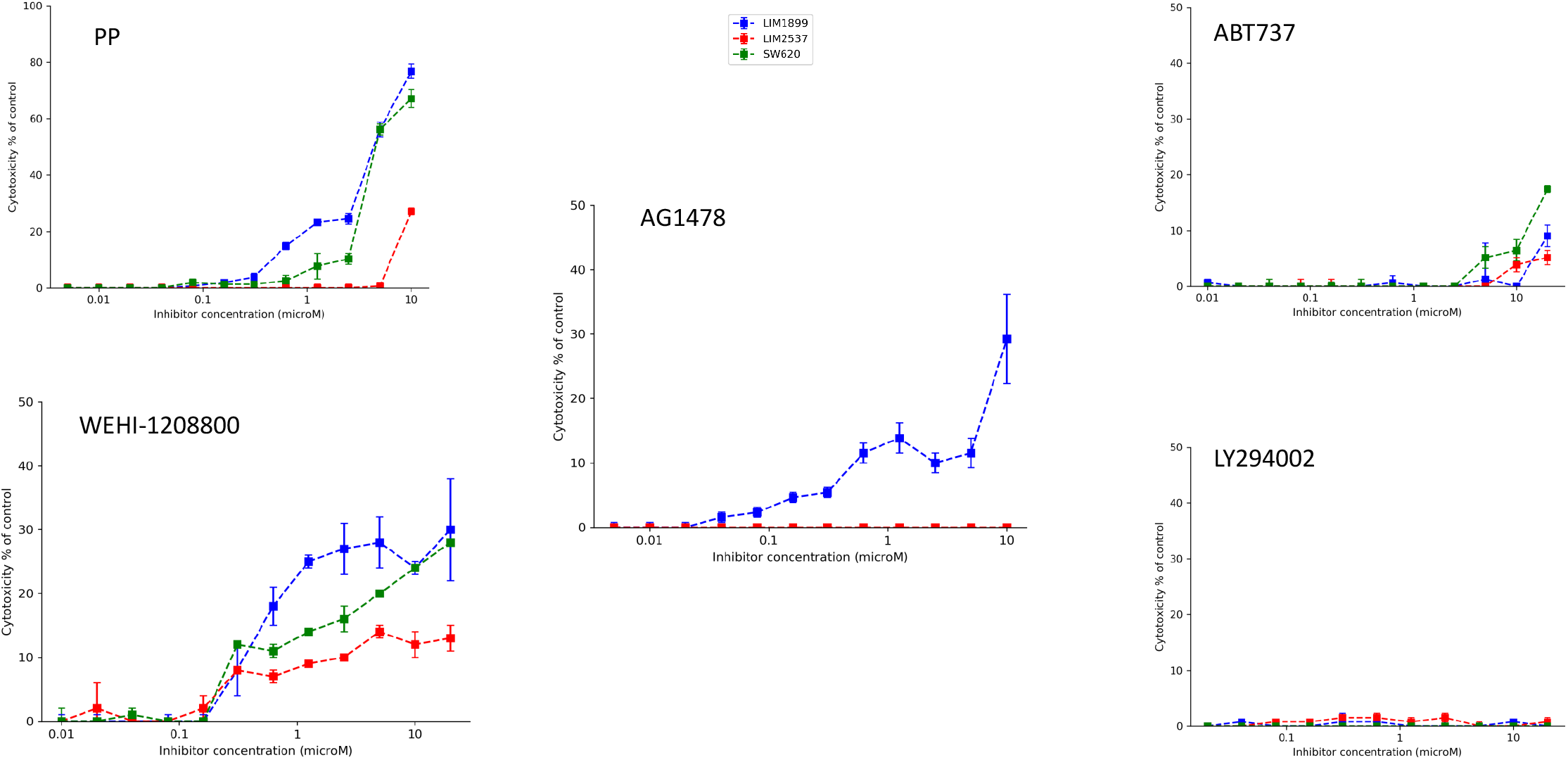
Adherent culture cytotoxicity of signaling inhibitors (pyrvinium pamoate PP, Wnt^i^; AG1478, EGFR^i^,; WEHI-1208800,src^i^;ABT737;bcl2^i^; and Ly294002,PI3K^i^) on three CRC cell lines (LIM1899,LIM2537 and SW620)

The PI3K inhibitor LY294002 partially inhibited the proliferation of LIM1899 at low concentrations and within the range already reported in the literature ^19^, whereas it was 4 times less potent in inhibiting LIM2537 cell line. As was the case for AG1478 (the EGFR small molecule inhibitor), the proliferation of SW620 cell line was resistant to inhibition of PI3K signaling pathway (IC_50_ ≥ 25μM) (Table 1 and Supplementary Figure 1).

At concentrations higher than 10μM, the BH3-only mimetic ABT737 inhibited LIM2537, LIM1899 and SW620 proliferation to 50% of control (Table 1 and Supplementary Figure 1). There are only a small number of studies on the effect of BH-3 only inhibitors on colorectal cancer cell lines *in vitro* 20, 21, but there is preliminary evidence of an increased response to chemotherapy when ABT737 is administered in a combination regimen for the treatment of other cancers ^22^. It must be noted that although ABT737 inhibits Bcl2, it also inhibits BclXL another member in this pro-survival family, so further studies with more specific inhibitors (e.g. ABT199 which targets Bcl2 more selectively^23^) would be needed to clarify if one or both of these proteins need to be inhibited to induce cytotoxicity.

As well as determining the IC_50_ for the effects of each compound on the cell lines, we assessed the maximum cytostatic effect of each drug on these adherent cell cultures (Supplementary Figure 3). All the compounds except DAPT induced considerable cytostasis (on at least two of the three cell lines) (Supplementary Figure 3). Although DAPT can sensitize colon cancer cell lines to cytotoxic agents in vitro ^24,25^, even 50μM DAPT did not inhibit proliferation the 3 cell lines and was therefore excluded from further testing.

Pyrvinium Pamoate is the only drug which completely abolished proliferation of all the 3 cell lines (Supplementary Figures 1,3). The SRC-inhibitor was also very effective in inducing cytostasis of the 3 cell lines: reducing proliferation of SW620 cells by 95%, of LIM2537 by 70% and of LIM1899 by 80%. LIM1899 cells were also 100% responsive to the cytostatic effect of the anti-EGFR small molecule inhibitor AG1478; whereas, as discussed above, SW620 cell line was resistant to EGFR inhibitor treatment (AG1478, Table 1) and the AG1478 induced only partial inhibition of proliferation in LIM2537 cells.

LIM1899 were the most sensitive cells to the cytostatic inhibition exerted by the PI3K inhibitor LY 294002, whereas SW620 showed only limited response to the inhibitor (Supplementary Figure 1). LIM2537, on the other hand, responded to LY 294002 with a 70% inhibition of proliferation. Finally, ABT737 displayed around 50% of maximum cytostatic effect on all 3 cell lines (Supplementary Figures 1,3).

### Cytotoxic Effects of signaling inhibitors

While the MTT assay assesses live cell numbers, it does not discriminate between inhibition of proliferation and induction of cell death. We analyzed the ability of the drugs used in the proliferation studies to induce death in SW620, LIM1899 and LIM2537 using the LDH Cytotoxicity Detection and compared the level of killing in cells cultured under both adherent and non-adherent (hanging drop) conditions (Figures 1 and 2, respectively).

Under adherent culture conditions AG1478 (10μM) only killed 30% of the LIM1899 (Figure 1) and AG1478 did not induce cell death in either LIM2537 cells or SW620 cells. The SRC-inhibitor WEHI1208800 was cytotoxic to SW620 cells (around 30% cytotoxicity at a concentration of 10μM), but only marginally cytotoxic for LIM1899 cells (around 10% at the same concentration) (Figure 1). This contrasts with its high potency as a cytostatic agent in the proliferation assays (Table 1 and Supplementary Figure 1). The EC_50_ for WEHI1208800 on SW620 is almost 4 times higher than the EC_50_ for the other two cell lines.

**Figure 2:**
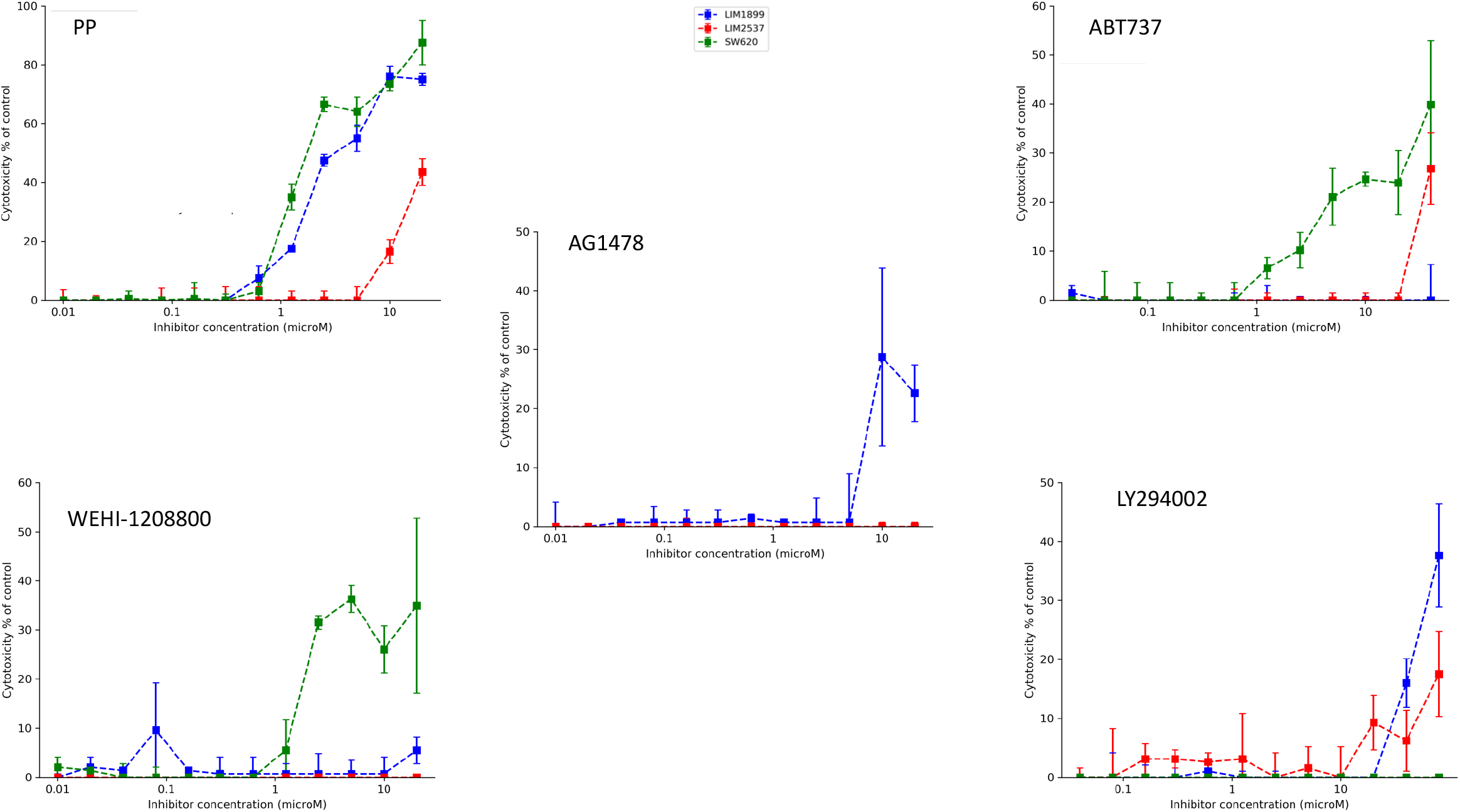
Non-Adherent culture cytotoxicity of signaling inhibitors (pyrvinium pamoate PP, Wnt^i^; AG1478, EGFR^i^; WEHI-1208800,Src^i^;ABT737;Bcl-2^i^; and Ly294002,PI3K^i^) on three CRC cell lines (LIM1899,LIM2537 and SW620)

Even at high concentrations (40μM), the PI3K inhibitor LY294002 does not exhibit any cytotoxic effect on adherent cultures of these cell lines and ABT737 exerts only marginal cytotoxic activity (Figure 1).

Pyrvinium Pamoate is by far the most potent cytotoxic drug in our panel, leading to 80% of cell death at 10μM on SW620 and LIM1899 cells (Figure 1); however it is not as potent for LIM2537 (maximum cytotoxic effect < 30%) and the EC_50_ is higher than for the other cell lines (8μM, see Table 1 and Figure 1). LIM2537 is the only heterozygous APC cell line amongst the three tested, so this result suggests a lower dependence for this cell line on Wnt signaling.

The CRC cell line cytotoxicity responses are different when cells are cultured in an anchorage-independent manner (Figure 2, Supplementary Figure 4; EC50’s are summarized in Supplementary Table 3). Within the same range of concentrations which are effective in adherent settings, SW620 is the only one of the three cell lines, sensitive to the SRC kinases inhibitor WEHI-1208800 (Figure 2); this drug reaches a maximum induced cell death of about 35-40% of control (Supplementary Figure 4). Interestingly, the PI3K-inhibitor LY294002, which did not exhibit any cytotoxic effect on any of the cell lines when cultured in monolayers, induced ∼ 40% cell death on the LIM1899 spheroids (Figure 2). The cytotoxic effect of LY294002 is exerted at higher concentrations (from 20µM onwards), however, the cytotoxic effect of LY294002 is only exerted at higher concentrations (from 20 µM onwards) where other cross-reactivities will be more common. These results may indicate a higher dependency on the PI3K pathways under non-adherent conditions, in accord with the observation that PI3K activation can abolish anoikis ^26^.

ABT737 was ineffective as a cytotoxic agent in monolayer cultures, but did show some cytotoxicity in non-adherent cultures of SW620 and LIM2537 (Figure 2). LIM1899 cell line remained unresponsive to the treatment with ABT737 (Figure 2). LIM1899 was the only cell line sensitive to the EGFR-inhibitor AG 1478 in the non-adherent cultures. There was a slight shift of the cytotoxicity curve to the right in comparison to the adherent cells; consequently, the EC50 values are higher (Supplementary Table 3). LIM2537 cells did not show any response to AG1478, mirroring the results obtained on the adherent cell cultures.

Pyrvinium Pamoate was again the most effective cytotoxic drug. Pyrvinium Pamoate caused around 70 to 85% of cell death in LIM1899 and SW620 (Figure 2 and Supplementary Figure 4). Although there was no change in the EC50 for PP on SW620 or LIM2537 cells, there was a significant shift to the left of the cytotoxicity curve for the LIM1899 cells in the non-adherent cultures (EC_50_ for cytotoxicity reduced from 3.5µM to 1.5µM) (Figure 2 and Supplementary Table 3).

### Sensitivity of colon cancer cells to combinations of targeting drugs in vitro

The drug combinations listed in Supplementary Table 4 were tested on two of the CRC cell lines, SW620 and LIM1899. The combination of the PI3K inhibitor LY 294002 and Wnt inhibitor PP significantly increased killing when compared to the single drug therapy in LIM1899 cells (Figure 3, Table 2).

**Table 2:** EC_50_ for inhibitors used as single agents and in combinations on CRC cell lines in 3D hanging drop cultures of colonospheres. EC_50_ are the average of triplicates from 3 experiments ± SD.

| Inhibitors (μM) | PP | PP<br>+ | PP<br>+ | PP<br>+ |  | WEHI-1208800<br>+ |  | AG1478<br>+ |  |  |
| --- | --- | --- | --- | --- | --- | --- | --- | --- | --- | --- |
| Cell lines |  | WEHI-1208800 | LY294002 | ABT-737 | WEHI-1208800 | ABT-737 | AG1478 | ABT-737 | LY294002 | ABT-737 |
| LIM1899 | 3 ± 0.1 | 2.7 ± 0.17 | 1.2 ± 0.06 | 2.6 ± 0.35 |  |  | >20 | >20 | >30 | >40 |
| SW620 | 6.5 ± 0.5 | 4.3 ± 1.5 | 1 ± 0.85 | 1.2 ± 0.7 | 4.5 ± 3.5 | 5 ± 2.8 |  |  | >40 | 6.6 ± 3.8 |

**Figure 3:**
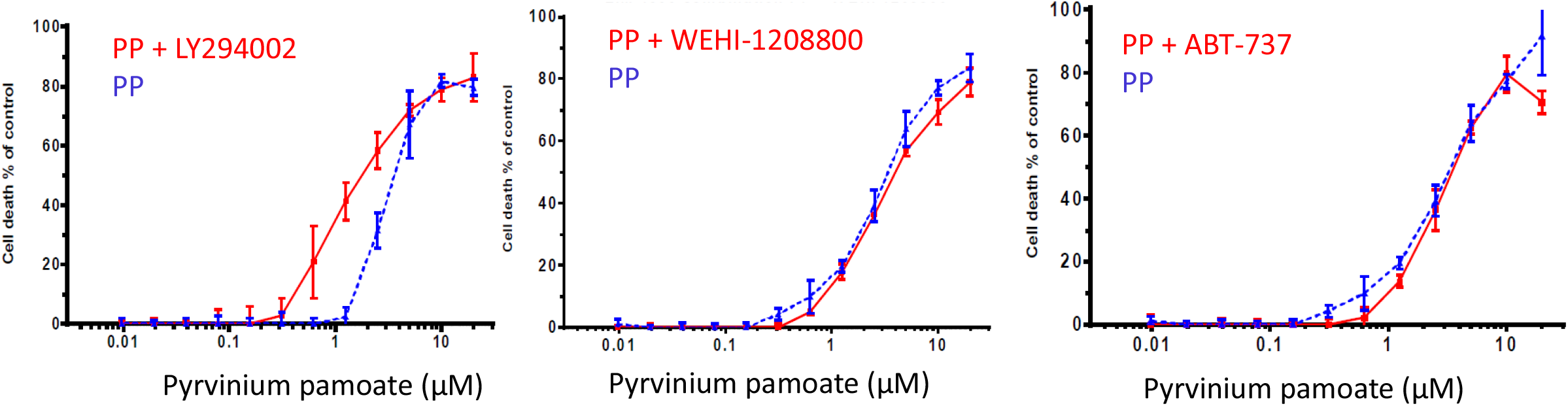
Cytotoxic effects of different concentrations of pyrvinium pamoate (PP) in combination with a fixed concentration of a PI3K inhibitor (LY294002,15μM), a Src inhibitor (WEHI-1208800, 1μM) or the pro-apoptotic drug (ABT737, 10μM) on the LIM1899 colorectal cancer cells growing as colonospheres in hanging drop cultures.

Neither the SRC inhibitor (WEHI-1208800) nor the Bcl family inhibitor (ABT737) elicited any increase in cell death or sensitivity to pyrvinium pamoate (Figure 3) in LIM1899 cells; nor did ABT737 induce any increase the sensitivity of SW620 cells to WEHI 1208800 (Supplemental Figure 5). Similarly, the SRC inhibitor (WEHI1208800) failed to increase the maximum cytotoxicity or the sensitivity of SW620 to pyrvinium pamoate (Supplementary Figure 5). However, pyrvinium pamoate in combination with either the PI3Kinase inhibitor (LY294002) or the Bcl-2 inhibitor (ABT737) elicits significant increases in sensitivity of SW620 cells to pyrvinium pamoate, i.e. a 6-fold decrease in the EC_50_ compared to pyrvinium pamoate as a single agent (Figure 4, Table 2). The shift to the left of the curve of cytotoxicity was more pronounced when Pyrvinium Pamoate was used together with ABT 737: it appears that ABT737 potentiates the cytotoxic effect of the Wnt inhibitor (pyrvinium pamoate, PP) starting from concentrations of PP as low as 30nM.

**Figure 4:**
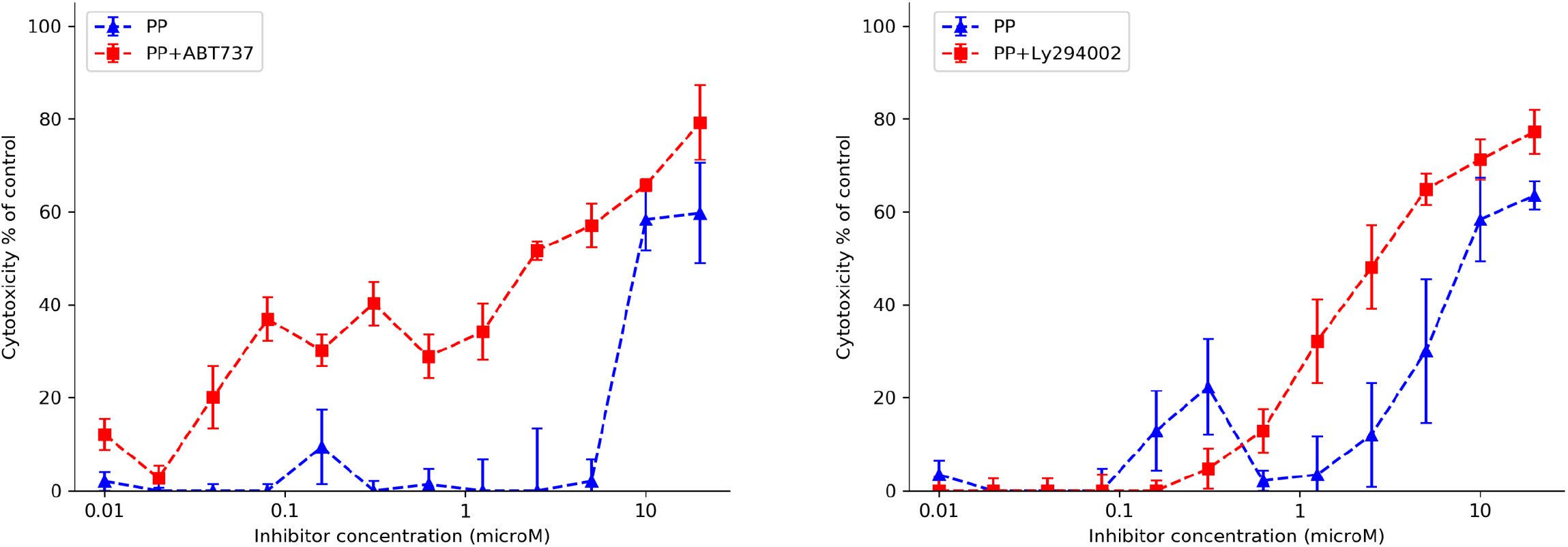
Cytotoxic effects of different concentrations of pyrvinium pamoate (PP) in combination with a fixed concentration of the pro-apoptotic drug (ABT737, 10μM) or a PI3K inhibitor (LY294002,15μM) on the SW620 colorectal cancer cells growing as colonospheres in hanging drop cultures.

From our initial results on the drug combinations studies, it was clear that the combination of the Wnt pathway inhibitor (Pyrvinium Pamoate, PP) and the Bcl-2 inhibitor (ABT737) killed SW620 effectively, with sensitization of the cells to PP at very low concentrations in comparison to when the drug was used as single agent (Figure 4). This same combination was ineffective on LIM1899 (Figure 3). SW620 carries an inactivating mutation of APC with loss of heterozygosity, whereas LIM1899 carries an activating β-catenin mutation. Moreover, SW620 cells are P53 mutant, whereas LIM1899 cells are P53 wild-type. These differences could be very relevant to the drug sensitivity, as PP is postulated to act by preventing β-catenin activation ^27,28^, while the loss of P53 function may upregulate the pro-life role Bcl-2 (Hemann,MT and Lowe SW, 2006).

Therefore, we investigated the hypothesis that this combination of inhibitors will be more effective on APC and/or P53 mutated cell lines, whereas it might not induce any cytotoxicity on β-catenin mutated/P53 wild type cell lines. To test this hypothesis, we examined more colorectal cancer cell lines, including 6 other lines carrying mutations of either the *APC* tumor suppressor gene or the *CTNNB1* (β-catenin) gene (see Supplementary Table 5). We treated these cell lines, growing as colonospheres in the hanging drop culture system, with the PP alone and in combination with ABT737 and assessed their responses using the LDH Cytotoxicity Assay. With the exception of LOVO colorectal cancer cell line, the APC mutant cell lines responded to the treatment with the PP and ABT737 combination with a shift in the EC_50_ towards the left of the cytotoxicity curve, i.e. increased sensitivity to PP (Figure 5). As expected, the PP/ABT737 combination induced apoptotic cell death as determined by FACS analysis (Supplementary Figure 6)

**Figure 5:**
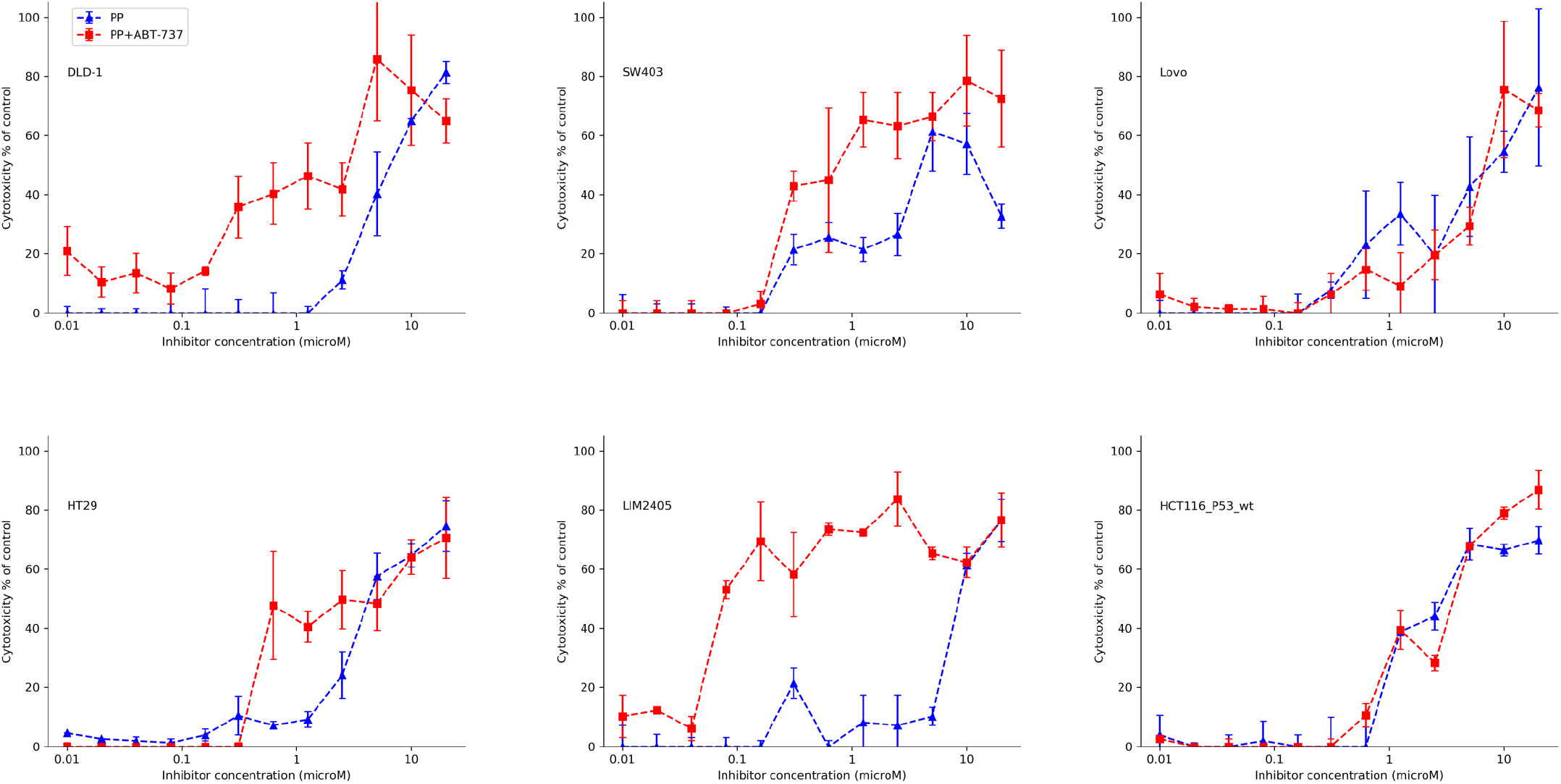
Cytotoxic effects of different concentrations of pyrvinium pamoate (PP) in combination with a fixed concentration of the pro-apoptotic drug (ABT737, 10μM) on six colorectal cancer cell lines (DLD-1,SW403,Lovo,HT29,LIM2405 and HCT116_P53_wt) growing as colonospheres in hanging drop cultures.

As we found for the β-catenin mutant cell line LIM1899 (see Figure 3), the HCT116 cells, which also have mutant β-catenin, were resistant to the combination of PP and ABT737 (Figure 5). LIM2405 and SW403 cells were 100-and 10-fold more sensitive to the combined treatment of PP plus ABT737 (Figure 5). As in the case of the other responsive cell line SW620, the addition of ABT737 potentiates the cytotoxic activity of PP at low concentrations.

### Effects of the pyrvinium pamoate/ABT737 drug combination on the growth of SW620 xenografts in mice

Many variables must be considered when switching from cell culture to small animals as experimental models; the process of translating the results obtained *in vitro* into informative experiments *in vivo* is often complex. However, two of the drugs used in our *in vitro* studies pyrvinium pamoate ^11^and ABT737 ^10^ have been used previously in mouse models of disease: ABT737 had been already used in mouse xenografts and reported to be quite effective in slowing the growth of subcutaneous tumors derived from SCLC cell lines (Oltersdorf et al 2005). ABT737 induced a complete regression of the tumors in this SCLC lung cancer model, when injected intra-peritoneum at 100 mg/kg/day, in 77% of the cases and significantly slowed tumor growth at a dose of 50 mg/kg/day ^11^. Pyrvinium Pamoate was used *in vivo* by Esumi and colleagues to treat subcutaneous human PANC-1 tumor xenografts in nude and NOD/SCID mice^10^. The drug was administered orally at a dose of 100 or 200μg/mouse/day and reached its maximal effect at 100μg/day.

Our aim in combining two drugs was to allow lower doses of each drug with equal or more tumor toxicity, thus minimizing unwanted side effects at the same time as achieving killing of the tumor cells. With this in mind, we decided to use ABT737 at a sub-maximal concentration of 50 mg/kg/day and pyrvinium pamoate at 50μg/kg/day. The two drugs were formulated for delivery in a mixture of 30% v/v propylene glycol, 5% v/v Tween 80, 65% D5W (5% w/v dextrose in water) and administered intra-peritoneum (I.P.) at 50mg/kg/day 5 days/week. Pyrvinium Pamoate, diluted in 2% v/v of DMSO in saline (final volume 200μl), was administered intragastrically to the mice by gavage at 50μg/mouse/day, 5 days/week. Administrations of the two drugs were separated by at least 3 hours gap to avoid cross reactions. Both drugs were also administered as single agents. Control mice received the vehicles with no drug. ABT737 is difficult to solubilize ^29^. Similarly, pyrvinium pamoate was not soluble in water, heating and stirring were required to keep the compound in solution before administration.

Treatment of the mice carrying the xenografts was started at day 7 after the SW620 cells were inoculated and terminated at day 35 after tumor inoculation. The remaining mice were euthanized and the tumors collected for histology. Spleen, livers and kidney from 4 mice per group were also collected for histology. The mice treated with the PP/ABT737 combination started to lose weight from day 10 onwards (Supplementary Figure 7A). We supplemented the mouse food with a combination of Sustagen and normal food at a ratio of 30:70, starting from day 13. Even with this addition, the mice treated with PP/ABT737 continued to lose weight. The average weight loss within the group in comparison to control was 15%, however, one mouse from the combination group needed to be euthanized for excessive weight loss (> 20%); 2 mice from the same group were euthanized after showing multiple signs of stress and sickness; 3 mice from the combination group died before the completion of the experiment. Unfortunately, the high general toxicity of the combination therapy, indicated by the body weight loss and the number of deaths in the combination group (75% of deaths within the group), together with the peritoneal precipitation of the ABT737 (Supplementary Figure 7B), indicated it was not safe to use ABT737 under the dose and administration regime chosen for this experiment. This finding was unexpected as it has not been reported in the literature before. Moreover an excessive weight loss may affect the overall metabolism of the mouse, with a catabolic status that would affect the growth of the tumors as well, when the tumor cells are deprived of the nutrients they need to keep multiplying. These facts heavily impacted on the overall significance of the *in vivo* model, underlining the need for finding good alternatives to the drug formulations and dosage currently in use, while maintaining the same inhibitory combination of targeted therapeutics (Wnt inhibitor and BH3-only mimetic) which achieved such promising results *in vitro*.

### Replacement of pyrvinium pamoate and ABT737 with orally available analogues

Fortunately oral analogues of both PP (pyrvinium phosphate, PPh^9^) and ABT737 (ABT263,^12^) were available. ABT263 has been shown to be highly effective in inducing regression of SCLC and ALL cell lines tumors in mouse xenografts as well as potentiating the anti-tumor activity of chemotherapeutic regimes already in use in B-cell malignancies (Tse et al 2008). PPh was synthesized from Pyrvinium Pamoate in our laboratory, following the process detailed by Yu and colleagues^9^. We tested the efficacy of PPh and ABT263 *in vitro* on SW620 cells grown in the hanging drop culture system. PPh and ABT263 were individually even more potent than PP and ABT737, respectively (Figure 6). We also tested the PPh/ABT263 combination on another CRC cell line (LIM2405) and again, the combination was more effective than the single agents at killing these CRC cells in the hanging drop cultures (Supplementary Figure 8).

**Figure 6:**
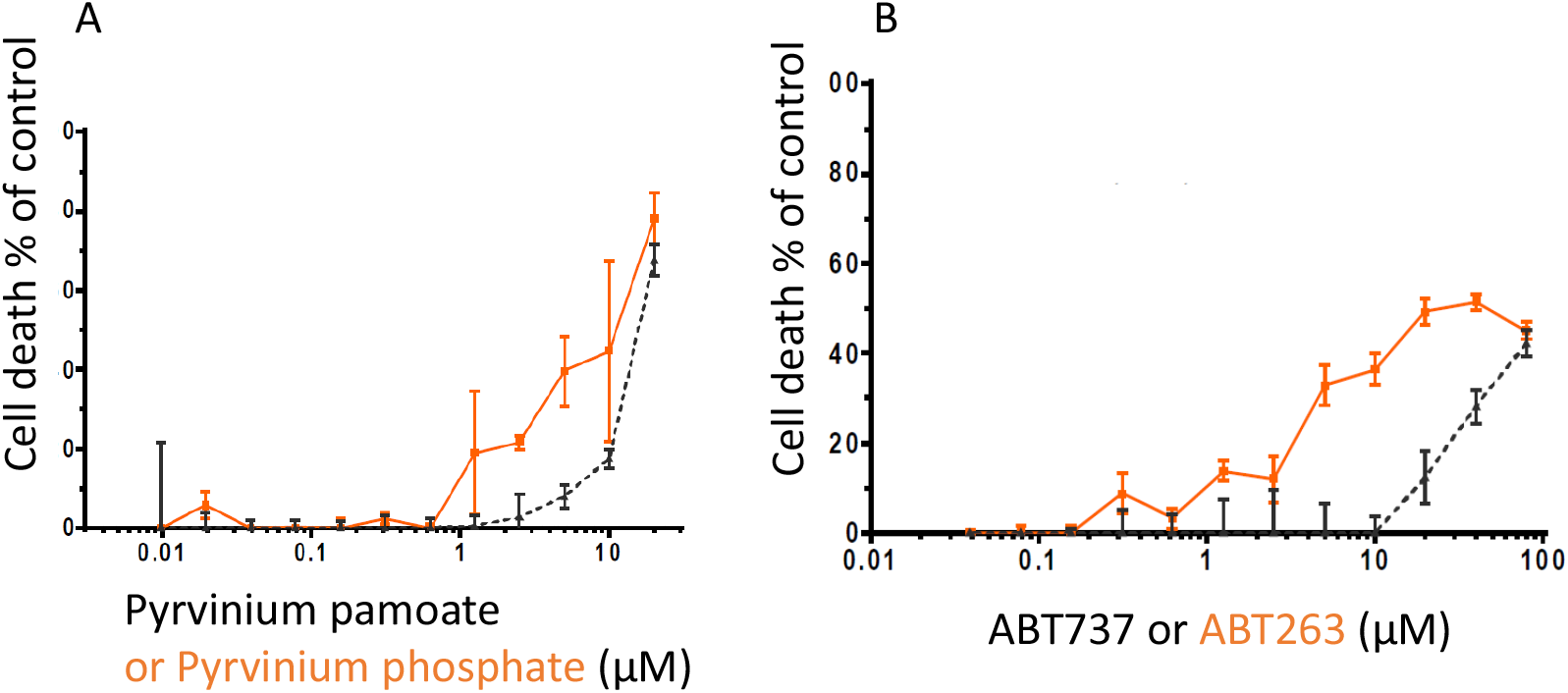
Cytotoxic effects of different concentrations of **A**, pyrvinium pamoate and pyrvinium phosphate; **B**, ABT737 and ABT263 on the SW620 CRC cell line growing as colonospheres in 3D-hanging drop cultures.

### Effects of the pyrvinium phosphate (PPh)/ABT263 treatment on the growth of SW620 xenografts in mice

The treatments with vehicle, PPh, ABT263 or the combined PPH/ABT263 were started 7 days after inoculating the tumor cells. The mice were inspected and weighed twice a week for the duration of the treatment. The tumor growth was measured with calipers whilst the mice were under a light general sedation. 50mg/kg/day of ABT263 diluted in a mixture of PolyEthylene Glycol (PEG) 30% v/v, Phosal 50 PG 60% v/v and Ethanol 10% was administered intra-gastrically by gavage (total volume of each dose was 200μl), 5 days/week. No problems were encountered in the preparation or administration of the ABT263 solution. 5mg/kg/day of Pyrvinium Phosphate (PPh) diluted in water was administered intra-gastrically by gavage in a maximum volume of 200μl for 5 days/week. To avoid direct interactions between the drugs and to allow the mice to recover from the gavage, the two drugs were administered separately (at least 3 hours apart). The experiment was repeated twice, the first time with NOD/SCID female mice, the second with NOD/SCID male mice. Each treatment group contained 8 to 10 mice. Treatment was continued for 23 days for the female mice and for 22 days for the male (Figure 7).

**Figure 7:**
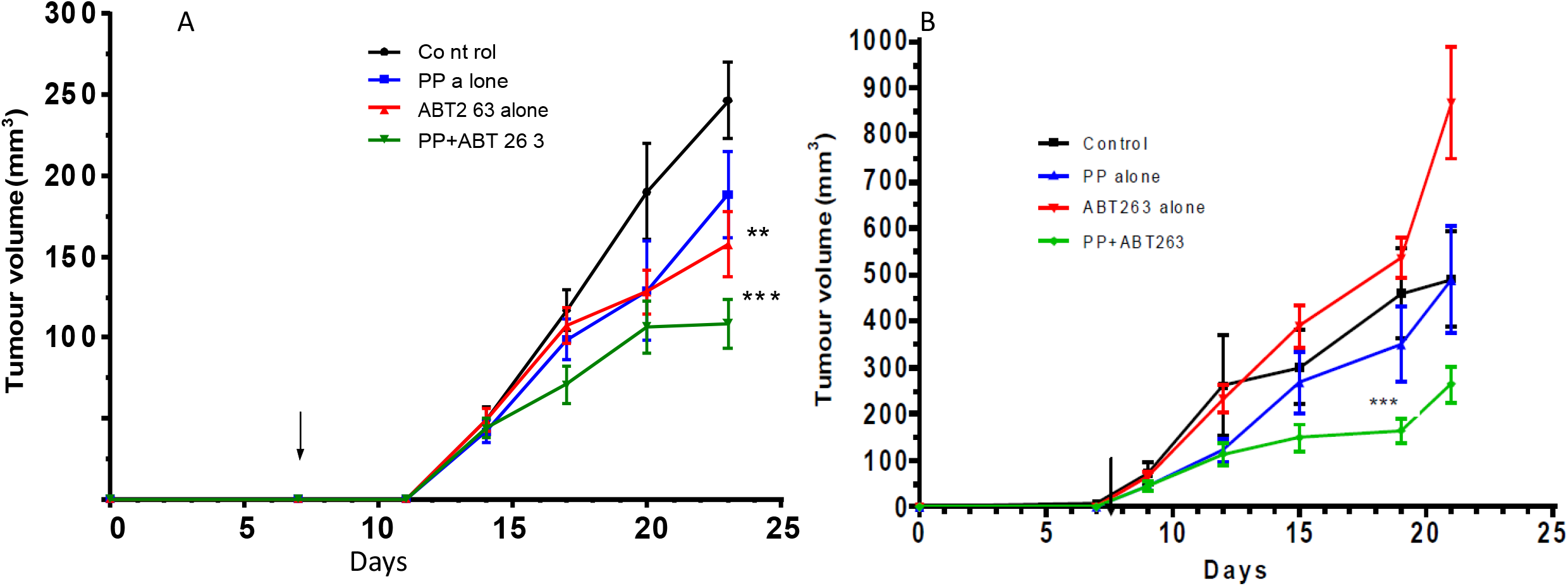
Effects of the Wnt inhibitor pyrvinium phosphate (PPh) and the pro-apoptotic Bcl-2 inhibitor ABT263 on the growth of SW620 tumor xenografts in **A**, female mice; **B**, male mice. The arrow shows the beginning of the treatment (day 7). The tumor volumes were measured twice a week, with the mice under general sedation. Each point represents the average of the volume of all the tumors within a group. Error bars = S.E.M., *** = p < 0.01, ** = p< 0.05.

Tumor xenografts from SW620 cells were established in both male and female cohorts and mice were treated with PPh and ABT263 alone and in combination (Figure 7). Female and male mice from the combination group and male mice treated with PPh as a single agent started showing body weight loss from day 10 and day 15, respectively (Supplementary Figure 9). Further weight loss was prevented by the dietary supplement (Sustagen). The average weight was less than 10% for both male and female mice, however, one mouse from the Pyrvinium Phosphate group was euthanized for excessive weight loss. We noted that the female cohort treated with PPh and ABT263 showed “matted fur” whereas mice treated with single agents did not. No other adverse health effects were observed.

Tumors in the PPh/ABT263 combination groups for both male and female cohorts exhibited slower growth rates in comparison to the controls and the single drug groups (Figure 7). In the male mice the tumor volume reached a plateau in the combination group on day 20. There was no difference between the growth of the SW620 tumors in the control group and the growth of the tumors in mice treated with PPh, however the ABT263 as a single agent decreased tumor growth (p<0.05) (Figure 7).

Histological analysis of the tumors showed that the vast majority were mucinous, with some tumors containing significant amounts of fluid. There were no signs of direct or indirect toxicity or damage to other organs and the tissue collected for histology looked normal by microscopy, with no detectable differences between treated and untreated samples as evidenced by H&E staining (Figure 8A-D).To assess whether the PPh/ABT263 treatment exerts *in vivo* anti-tumor activity through induction of apoptosis, the levels of apoptosis in SW620 xenograft tumors were assessed using immunohistochemical staining with ApopTag^30,31^. To exclude stromal tissue from the analysis, we stained two consecutive sections for each tumor, one with hematoxylin eosin, which allows a clear distinction between stroma and glandular epithelium, and the other for apoptosis (Figure 8A-D). The percentage of apoptotic cells was close to zero in tumor sections derived from untreated and PPh treated mice (Figure 8A, B). ABT263 treated tumors displayed only a slightly higher number of apoptotic cells in comparison to the control and PPh treated mice (Figure 8C). Whilst the frequency of apoptic cells appeared to be higher in tissue from the PPh/ABT263 combination (Figures 8D, E), there was no statistical difference when compared to the single agent treated or the control mice (p<0.25).

**Figure 8:**
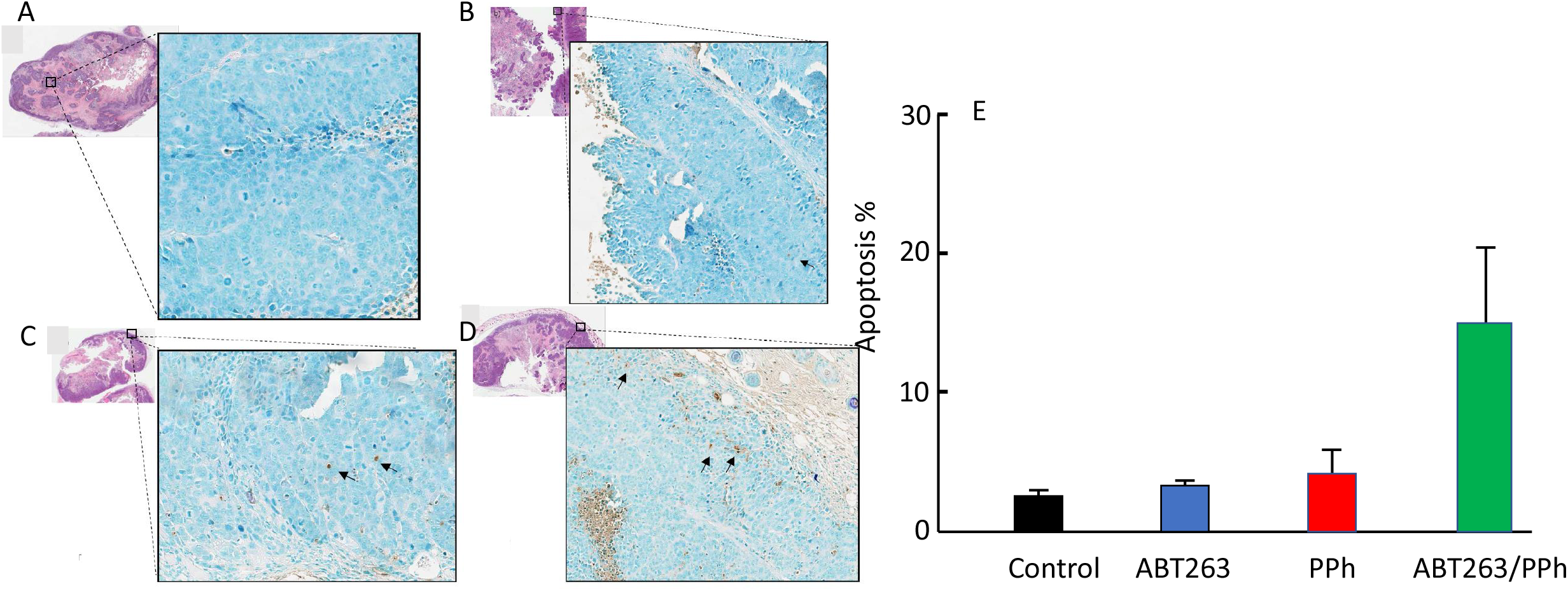
Detecting apoptotic cells (ApopTag IHC) in the SW620 tumor xenografts treated with **A**, Vehicle, **B**, Pyrvinium phosphate (PPh), **C**, ABT263 or **D**, PPh +ABT263. A total of 64 sections were stained using the Apoptag peroxidase Apoptosis Detection Kit (2 sections/tumor, 2 tumors/mouse, 4 mice/treatment group). 20 fields per section were chosen at random within the glandular epithelium (excluding stromal component and necrotic/infiltrating inflammatory tissue). The apoptotic cells (brown) and the total number of cells were counted within each field and the percentage of apoptosis calculated per field. The average percentage of apoptosis was then calculated for each section and the results averaged per treatment group. The graph shows the average total percentage of apoptosis in treatment each group expressed as mean ± SD. The apparent increase in apoptosis induced by the combination treatment was not statistically significant in comparison to control (P = 0.25). Metamorph and GraphPad were used to perform the statistical analysis of the results: an unpaired t-test was performed to calculate the P values.

## Discussion

The colorectal cancer cell lines we tested respond to one or more single agent targeted treatments in both adherent cell cultures and the non-adherent hanging-drop culture system. However, in the clinic, colon cancer patients^32^ receiving targeted therapies usually relapse and die as a result of their cancer^33^. Although the adherent cell proliferation was blocked by the EGFR, Wnt and src inhibitors, the PI3K inhibitor was only active in the non-adherent cultures on two of the cell lines and the src inhibitor was only active on two of the three cell lines in the non-adherent cultures. This suggests that integrin signaling is likely to modify responses to anti-cancer targeting drugs. Some consideration should be given to modulating integrin signaling ^34,35^ when trying to optimise the use of anti-cancer drugs for treating advanced cancers. One of our cell lines (LIM1899) was resistant to the pro-apoptotic drug ABT737. Now that there are several options for inhibiting the pro-survival pathways, other BH3 inhibitors should be tested for activity on colorectal cancer cell ^20,36^.

In our experiments PP was the most consistent and potent cytotoxic agent for the three cell lines, suggesting that inhibition of Wnt signaling should be a key component option for treating advanced cancers. In five CRC cell lines with APC mutations, treatment with PP was enhanced by the presence of the pro-apoptotic agent ABT737. For example, SW620 cells are 6-fold more sensitive to PP in the presence of ABT737 or the PI3K inhibitor (LY294002)^37^. In cells with *CTNNB*1 mutations, the cells were killed by PP, but there was no increased potency in the presence of the pro-apoptotic drug. Our results suggest, genetic screening^38^ and or profiling for the expression levels of the Bcl-2 family members^32^ will be helpful in predicting which patients might be the best responders to the dual Wnt inhibitor/pro-apoptotic drug treatment.

Despite successful reports which use either ABT737 ^39^ or PP mouse models^40^, we found the formulation and precipitation in the peritoneum renders these agents unsuitable for use *in vivo*. Fortunately, soluble analogues of both ABT737 (ABT263, ^12^) and pyrvinium phosphate (PPh, ^9^) were readily available and appeared to work just as well *in vitro*. By themselves neither PPh nor ABT263 reduced SW620 tumor growth, however, in combination, PPh with ABT263 showed low toxicity and was effective for reducing the growth of SW620 tumor xenografts. Where patients are suffering with advanced colorectal cancers and carrying *APC* mutations consideration should be given to dual drug treatments which target Wnt signaling and enhance apoptosis. It would be interesting to know which of the pro-apoptotic drugs is the best sensitizer for PPh and whether more specific/potent Wnt inhibitors might be even more effective. Both PPh and ABT263 can be administered orally (as in our experiments), but other routes of administrations should be tested. Given the increased potency of PPh in the presence of a pro-apoptotic drug, it is conceivable that a dual drug treatment (Wnt inhibitor plus a pro-apoptotic drug) will also kill colon adenoma stem cells (most of which have APC mutations ^41^) and thus act as a chemoprevention strategy for reducing the incidence of colorectal cancer. Consequently, we have extended our studies to investigate the effects of PPh, ABT263 and anti-inflammatory drug combinations on two models of CRC initiation: APC^min^ and DClk1-Cre mice primed with DSS (Dextran, sodium sulfate)^42^.

## Acknowledgements

The authors acknowledge support from the Ludwig Institute for Cancer Research, WEHI and the National Health and Medical Research Council (NH&MRC): Program Grant #487922 and Development Grant #1017059. The funding body had no role in the design of the study, collection, analysis or interpretation of data or in writing the manuscript.

## Authors’ Contributions

SPC, FW, JW performed the experiments. SPC, FW, GL and AWB guided the design and interpretation of the experiments and wrote the first drafts of the manuscript. SPC, FW, MCF and AWB analysed and interpreted the data and contributed to the critical evaluation of the manuscript. All authors contributed to the final text of the manuscript.

