## Supplemental_Information for "Dual Drug Targeting to Kill Colon Cancer Cells"

**Corona et al**

**Supplementary Information**

**Supplementary Methods**

**TCF-GFP Reporter assay** The TCF-GFP Cignal Reporter assay (Qiagen, Hamburg, Germany) allows quantitation of the activation of Wnt dependent transcription in cultured cells^1, 2^. This assay is based on the TCF/LEF Luciferase assay, but the fluorescent protein is the GFP (Green Fluorescent Protein reporter gene). The TCF/LEF is linked to the GFP through a TATA box promoter so that when Wnt target genes transcription is activated, the cells produce GFP which can be measured by immunofluorescence or by FACS analysis.

**Cell transfection with Fugene HD** Transfection of colon cancer cell lines with the TCF-GFP Cignal Reporter assay was carried out using Fugene HD transfection reagent (Promega, Madison, WI), following the manufacturer’s instructions. Briefly, SW 620 and LIM 1899 colon cancer cells were plated at a density of 5 x 10^4^/well in 96 well/plates. After 24 hours, once the cells had attached to the plastic substrate, Fugene HD was diluted into serum-free culture medium in a sterile Eppendorf tube, at concentration of 6μl in 500μl of medium (the optimized concentration for each cell line was chosen after testing serial ratios in trials experiments). The Reporter, positive control and negative control cDNA was diluted 1:25 in serum-free medium into another sterile Eppendorf tube at a final concentration of 4μg/ml (20μl of cDNA in 500μl of medium). After 5 minutes incubation at room temperature, the Fugene HD solution was added to the cDNA drop-wise and this solution incubated at room temperature for 15 minutes. The medium was then replaced with fresh cell culture medium in the wells containing the cells and the reagent mix added to the medium at a 1:1 ratio (final volume 100μl/well).24h later the medium was diluted 1:2 using normal medium plus 10% Fetal Calf Serum. Transfection efficiency was monitored via GFP expression at 24 and 48hrs using fluorescence microscopy

**Assessment of Pyrvinium Pamoate activity on TCF-GFP transcription** 48h after transfection pyrvinium pamoate was added to a final concentration of 2μM to the cells, after confirming the expression of GFP in the positive control wells by fluorescence microscopy. Cells were than incubated for a further 18h or 29h with or without the drug before imaging each well. Images for the TCF/LEF Reporter Assay (GFP) were taken on a Nikon C1 Confocal Microscope after adding Hoechst 33342 live staining dye (Sigma Aldrich) at a final concentration of 25μM in culture medium, 30 minutes before taking pictures, to highlight the nuclei, thus allowing cell counting. Images were taken with the bright-field, the green fluorescence channel for the GFP-positive cells and the blue fluorescence channel for the nuclei, with a Nikon Plan Fluor lens, at 10x magnification.

MetaMorph version 7.7.10 was used to count the percentage of green cells over the total number of cells per field. GraphPad Prism 6 was used for the graphs.

**Supplementary Figure Legends**

**Supplementary Figure 1** : The effects of different concentrations of signaling inhibitors (AG1478, EGFR^i^; ABT737, Bcl2^i^; WEHI1208800, Src^i^; DAPT, notch^i^; LY294002, PI3K^i^ and Pyrvinium pamoate, Wnt^i^) on the proliferation of three CRC cell lines (LIM1899,LIM2537 and SW620) in adherent cell cultures, as measured using the MTT assay. The best fit of the inhibition of proliferation curve is shown. Error bars are mean ± SEM from three experiments.

**Supplemental Figure 2 :** TCF-GFP Reporter assay indicates that pyrvinium pamoate inhibits Wnt signaling in LIM1899 and SW620 cells. SW 620 and LIM 1899 cells were plated at a density of 5 x 10^4^ in 100μl of normal medium/well in 96 well/plates. Cells were transfected with 0.75μl of Fugene HD and 1μl of vector DNA (positive, negative or TCF-GFP) in 25μl of Optimem transfection medium. Both controls (positive and negative) were plated in duplicates, whereas the TCF-reporter was plated in triplicates. 24 hours after transfection PP was added to the cells at a concentration of 2μM. Cells were incubated for further 32h and imaged with a fluorescent microscope at two time-points (18h and 29h). The pictures were taken at the 18h time point unless otherwise specified.

No fluorescence is evident in the negative controls, both with and without the drug. The number of fluorescent cells per field is not affected by the drug in the cells transfected with positive control. Cells transfected with the TCF-GFP vector show a consistent decrease in fluorescence when exposed to the drug for 18 and 29h. Images were taken using a Nikon C1 Confocal Microscope (10x) under fluorescent light.

**Supplementary Figure 3** : The maximum proliferative inhibitions for each signaling inhibitor on three CRC cell lines (see details legend Supplementary Figure 1) are plotted as the mean of three experiments with triplicate assays for each experiment. Apart from the lack of effect of DAPT on the LIM1899 and SW620 cultures, all the results showed statistically significant inhibition in comparison to the control cultures (i.e. P<0.005).

**Supplementary Figure 4** : Maximum cytotoxic effect of drugs on cells cultured as spheroids in Hanging Drops. Quantitation of the maximum percentage of apoptotic cell-death in comparison to controls, measured using the Cytotoxicity Detection Assay by Roche. Pyrvinium Pamoate induces 70% of cell death in SW 620 and LIM 1899 cell lines. All the other drugs are only moderately cytotoxic. LY294002 does not induce any cell death under these culture conditions. AG1478 is not cytotoxic on LIM 2537, whilst it was not tested with this assay on SW 620 because of the absence of EGFR on these cells. The graph shows the average results from 3 experiments, each experiment performed with triplicate wells. Data are drawn as mean ± SEM.

**Supplementary Figure 5** : A, WEHI1208800 and ABT737 (10μM) do not synergize when used in combination on SW620 cells; B, Pyrvinium pamoate and WEHI1208800 (1μm) do not synergize when used in combination in the cytotoxicity assay with SW620 cells.

**Supplementary Figure 6** : Treatment of SW620 cells with PP/ABT737 induces cell death via apoptosis. Cells were cultured as colonospheres in hanging drops and exposed to vehicle only or to Pyrvinium Pamoate and ABT 737 as single agents or in combination (PP 1μM and ABT 737 10μM). After 24h the colonospheres were disrupted into a single cell suspension and fixed in methanol. Cells were incubated with RNAse and stained with Propidium Iodide prior to analysis by FACS. At least 10,000 cells were analyzed for each sample. The upper panel shows the area chosen for the analysis, with exclusion of the dead cells. The purple histogram is the control sample. SW 620 cell line showed a significant increase in the percentage of apoptotic cells visible in the sub-G1 section (M1) of the graph when treated with the combination in comparison to single agents and untreated cells.

**Supplementary Figure 7 :** A, Effect of the PP and ABT 737 combination treatment on mice body weight. Number of mice shown for every day of the experiment. The upper part of the graph shows the average weight of the mice (left axis) in each group throughout the duration of the experiment. The mice from the combination group lost significant weight starting from day 10 onwards and never regained the original weight, even after supplementation with Sustagen (started on day 16th, arrow). The mice from the ABT 737 single agent group showed a modest body weight loss, in comparison to controls, more accentuated towards the end of the experiment. Each point represents the average of all the mice weights within each group. Error bars are S.E.M. The lower part of the graph summarizes the number of mice (right axis) left in each group each day. In the last week of therapy, we were left with 3 and ultimately 2 mice within the combination group, due to the high general toxicity of the combination under the chosen conditions of administration. To be noted that the control group started with 1 mouse less than the other groups. One mouse from the Control group showed signs of distress and sickness and needed to be euthanized on day 27; B, PP and ABT 737 combination group mouse autopsy, xenograft trial. At the end of the experiment, 4 mice randomly selected within each group were subjected to autopsy and inspection of the peritoneal cavity and organs. At the inspection of the mice from the combination therapy group, a yellow, viscous precipitate was visible, covering all the internal surface of the peritoneum and the contained organs. Liver, small and large intestine were heavily involved with the intestine forming adhesions and with other signs of peritoneal inflammation. The arrow shows an area of ulceration of the visceral peritoneum attached to the small intestine, with involvement of the intestinal wall.

**Supplementary Figure 8** : Cytotoxicty effects of treatment with different concentrations of purvinium phosphate (PPh), ABT263 and the PPh/ABT263 (0.3μM) combination on the CRC cell line LIM2405 growing in 3D-hanging drop cultures.

**Supplementary Figure 9 :** A, Effect on Body Weight of treating **female** mice carrying SW620 xenografts with Pyrvinium phosphate (PPh) and ABT263. The mice were weighed twice each week. The combination group appeared to lose weight from day 8 to 10. The food for all of the mice was then supplemented with Sustagen; B Effect on Body Weight of treating **male** mice carrying SW620 xenografts with Pyrvinium phosphate (PPh)and ABT263. The mice were weighed twice a week. Two groups (PP and combination) appeared to lose weight from day 15 The food for all of the mice was then supplemented with Sustagen.

**List of Supplementary Tables**

**Supplementary Table 1** : Mutations in colorectal cell lines analyzed for drug sensitivity

**Supplementary Table 2** : Summary of Signaling inhibitors used in this report to treat CRC cell lines

**Supplementary Table 3** : Summary of EC50s for the cytotoxicity effects of the compounds used to treat each cell line in cells grown in 3-D hanging drop cultures. Each experiment was done in triplicate, mean ± SD is shown. WEHI = WEHI 1208800; PP = Pyrvinium Pamoate

**Supplementary Table 4** : Summary of inhibitor combinations used to treat the three CRC cell lines

(SW620, LIM2537 and LIM1899) in 3D hanging drop cultures.

**Supplementary Table 5** : Summary of Oncogene and Tumour Suppressor Gene profiles for CRC cell lines^3^: DLD-1^4, 5^, LOVO ^4, 5^, HT-29^4, 5^, LIM 2405^4-6^: SW 403 ^4, 5^ and HCT116 ^4, 5^

* HCT116 parental colorectal cancer cell line and its derivative HCT116p53KO which has two inactivated p53 alleles^7^ were tested :
