## Supplemental_Tables_Figs for "Dual Drug Targeting to Kill Colon Cancer Cells"

Corona et al

Supplementary Tables and Figures

Supplementary Table 1 Mutations in colorectal cell lines analyzed for drug sensitivity

| Mutations Cell line | APC | β-catenin | K-Ras | B-Raf ex15 | PI-3-K | p53 |
|---|---|---|---|---|---|---|
| LIM2537 | wt, stop1472 | wt | wt | V600E | Δ exon13 (p85) | G245D |
| LIM1899 | Wt | S45P | wt, G12A | wt | wt | wt |
| SW620 | stop1338 | wt | G12V | wt | wt | R273H + P309S |

Supplementary Table 2 Summary of Signaling inhibitors used in this report to treat CRC cell lines

| Drug | Mechanism of action | Bibliography |
|---|---|---|
| AG1478 | EGFR inhibitor<br>Competitive inhibition of the ATP binding site of the kinase | Ellis *et al.*,<br>2006 |
| WEHI 1208800 | SRC Family Inhibitor<br>Competitive inhibition of the ATP binding site of the kinase | Patent No.<br>PCT/AU2011/000858,<br>8/07/2011 |
| LY294002 | PI3K inhibitor<br>Competitive inhibition of the ATP binding site of the kinase | Vlahos *et al.*,1994 |
| ABT-737<br>ABT-263 | Bcl-2 Family Inhibitor<br>Allosteric inhibition of the α-helix region (BH3) | Oltersdorf *et al.*, 2005 |
| Pyrvinium Pamoate<br>Pyrvinium Phosphate | WNT Inhibitor<br>Activation of GSK-3β | Venerando et al., 2013 |
| DAPT | NOTCH signaling inhibitor, γ-secretase inhibitor<br>Competitive inhibition of Preselenin | Morohashi *et al.*, 2006 |

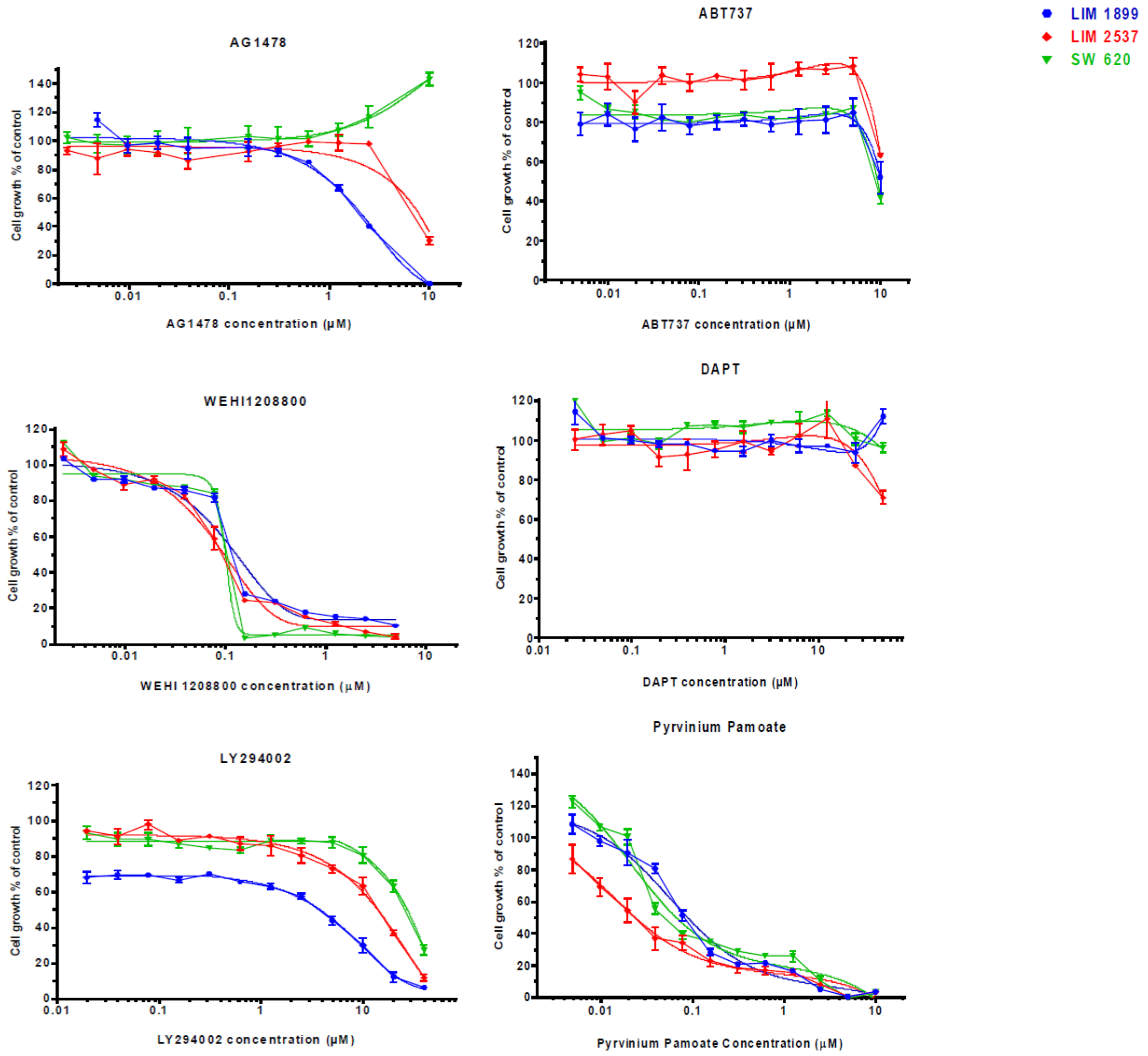


**Supplementary Figure 1** The effects of different concentrations of signaling inhibitors (AG1478, EGFR$^i$; ABT737, Bcl2$^i$; WEHI1208800, Src$^i$; DAPT, Notch$^i$; LY294002, PI3K$^i$ and Pyrvinium pamoate, Wnt$^i$ )on the proliferation of three CRC cell lines (LIM1899,LIM2537 and SW620) in adherent cell cultures, as measured using the MTT assay.

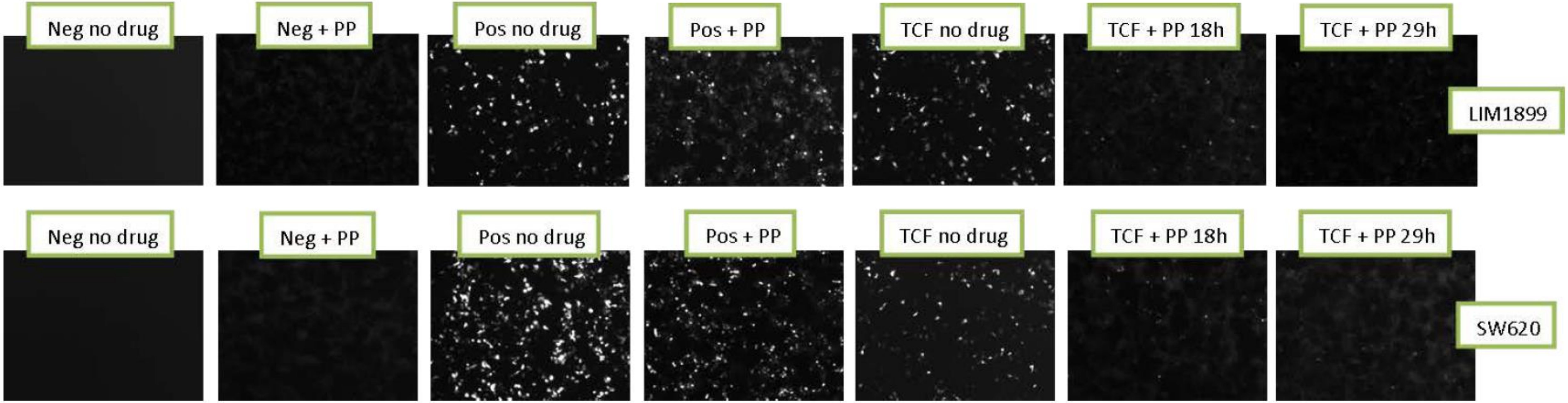


**Supplemental Figure 2 :** TCF-GFP Reporter assay indicates that pyrvinium pamoate inhibits Wnt signaling in LIM1899 and SW620 cells. SW 620 and LIM 1899 cells were plated at a density of 5 x $10^4$ in 100µl of normal medium/well in 96 well/plates. Cells were transfected with 0.75µl of Fugene HD and 1µl of vector DNA (positive, negative or TCF-GFP) in 25µl of Optimem transfection medium. Both controls (positive and negative) were plated in duplicates, whereas the TCF-reporter was plated in triplicates. 24 hours after transfection PP was added to the cells at a concentration of 2µM. Cells were incubated for further 32h and imaged with a fluorescent microscope at two time-points (18h and 29h). The pictures were taken at the 18h time point unless otherwise specified. No fluorescence is evident in the negative controls, both with and without the drug. The number of fluorescent cells per field is not affected by the drug in the cells transfected with positive control. Cells transfected with the TCF-GFP vector show a consistent decrease in fluorescence when exposed to the drug for 18 and 29h. Images were taken using a Nikon C1 Confocal Microscope (10x) under fluorescent light.

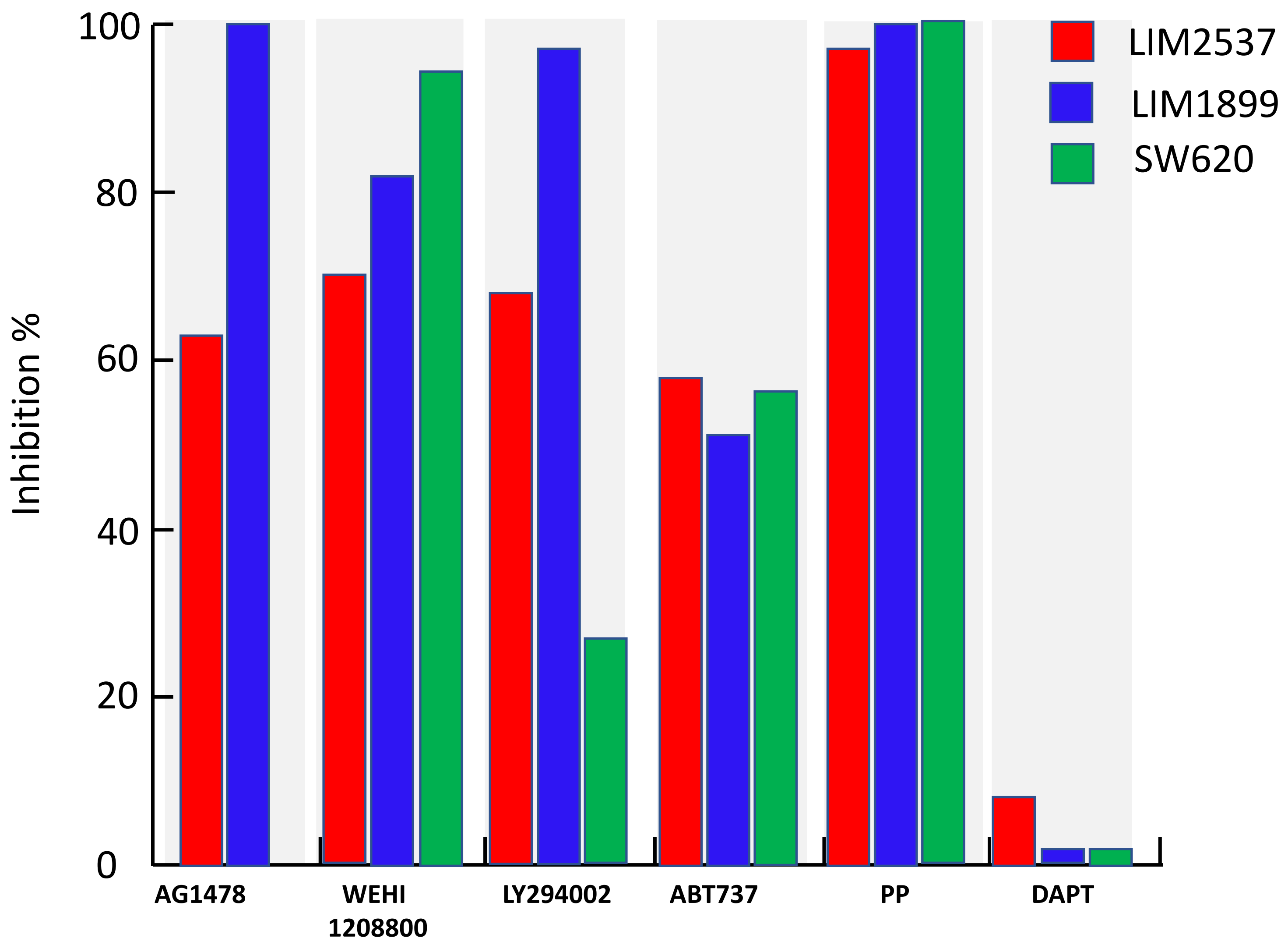


**Supplementary Figure 3** The maximum proliferative inhibitions induced by each of the signaling inhibitors on three CRC cell lines in adherent cell assays (see details legend Supplementary Figure 1) are plotted as the mean of three experiments with triplicate assays for each experiment. Apart from the lack of effect of DAPT on the LIM1899 and SW620 cultures, all the results showed statistically significant inhibition in comparison to the

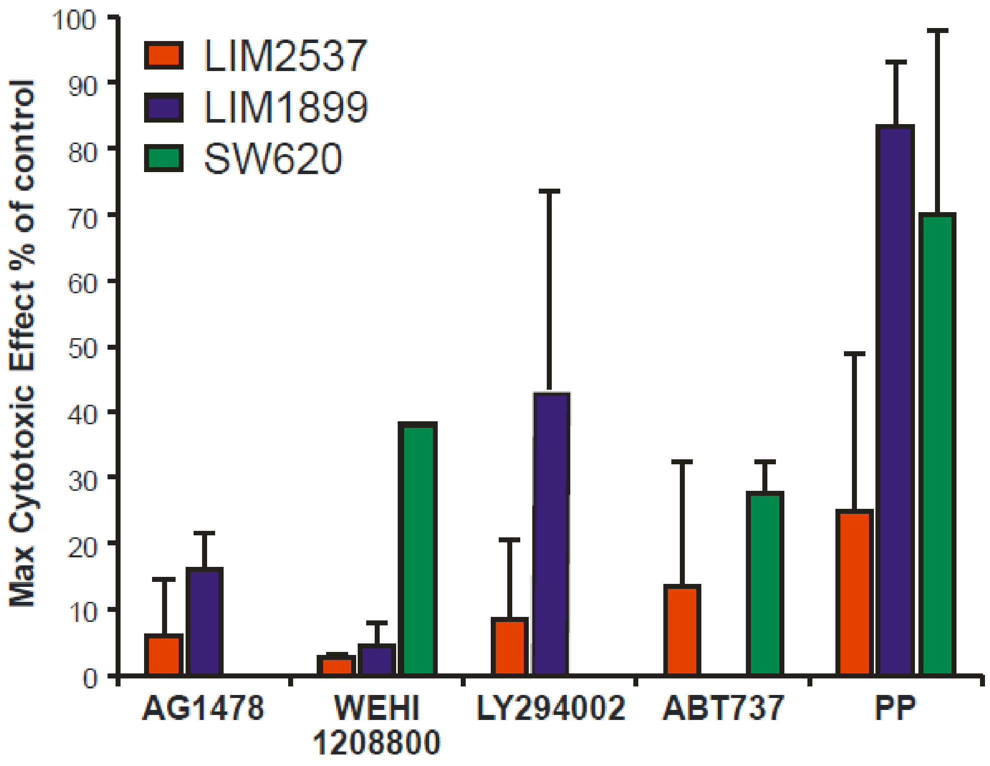


**Supplementary Figure 4 Maximum cytotoxic effect of drugs on cells cultured as spheroids in Hanging Drops.** Quantitation of the maximum percentage of apoptotic cell-death in comparison to controls, measured using the Cytotoxicity Detection Assay by Roche. Pyrvinium Pamoate induces 70% of cell death in SW 620 and LIM 1899 cell lines. All the other drugs are only moderately cytotoxic. LY294002 does not induce any cell death under these culture conditions. AG1478 is not cytotoxic on LIM 2537, whilst it was not tested with this assay on SW 620 because of the absence of EGFR on these cells. The graph shows the average results from 3 experiments, each experiment performed with triplicate wells. Data are drawn as mean ± SEM.

Supplementary Table 3: EC50s for cytotoxicity of the compounds for each cell line in cells grown in 3-D culture. Each experiment was done in triplicate, mean ± SD is shown.
WEHI = WEHI 1208800; PP = Pyrvinium Pamoate

| Inhibitors (µM) $EC_{50}$ | AG1478 | WEHI | LY294002 | ABT737 | PP |
|---|---|---|---|---|---|
| LIM2537 | 6 ± 8.4 | 0.6 ± 0.1 | 23 ± 32.5 | 11 ± 15 | 14 ± 5.6 |
| LIM1899 | 11 ± 3 | 8.6 ± 8 | 35 ± 18 | >40 | 1.6 ± 0.9 |
| SW620 | | 0.9 ± 0.8 | >80 | 6.6 ± 3.8 | 6 ± 2.3 |

Supplementary Table 4 Summary of inhibitor combinations tested on the three CRC cell lines: SW620, LIM2537 and LIM1899

| Cell lines | Small Molecule Inhibitor Combinations |
|---|---|
| SW 620 | PP + WEHI1208800<br>PP + ABT 737<br>PP + LY294002<br><br>WEHI1208800 + ABT 737 |
| LIM 1899 | PP + WEHI1208800<br>PP + ABT 737<br>PP + LY294002<br><br>AG 1478 + ABT 737 |
| LIM 2537 | WEHI1208800 + ABT 737 |

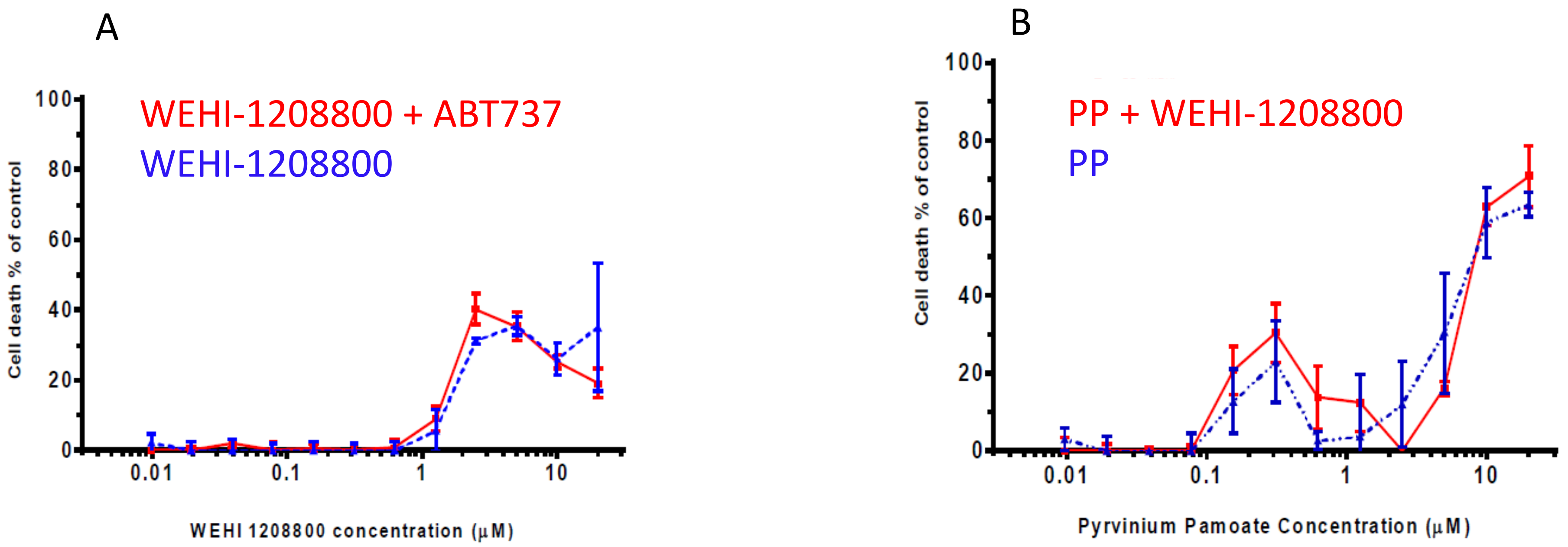


**Supplementary Figure 5** A, WEHI1208800 and ABT737 (10µM) do not synergize when used in combination on SW620 cells; B, Pyrvinium pamoate and WEHI1208800 (1µm) do not synergize when used in combination on SW620 cells

Supplementary Table 5 Summary of Oncogene and Tumour Suppressor Gene profiles for CRC cell lines used in this study

| Mutations / Cell line | APC | B-Catenin | K-Ras | B-Raf | PI3K | P53 |
|---|---|---|---|---|---|---|
| DLD-1 | *mut* | *wt* | *mut* | *wt* | *mut* | *mut* |
| LOVO | *mut* | *wt* | *mut* | *wt* | *wt* | *wt* |
| HT-29 | *mut* | *wt* | *wt* | *mut* | *mut* | *mut* |
| LIM2405 | *mut* | *wt* | *wt* | *mut* | *wt* | *wt* |
| SW403 | *mut* | *wt* | *mut* | *wt* | *wt* | *wt* |
| HCT116 | *wt* | *mut* | *mut* | *wt* | *mut* | *wt** |

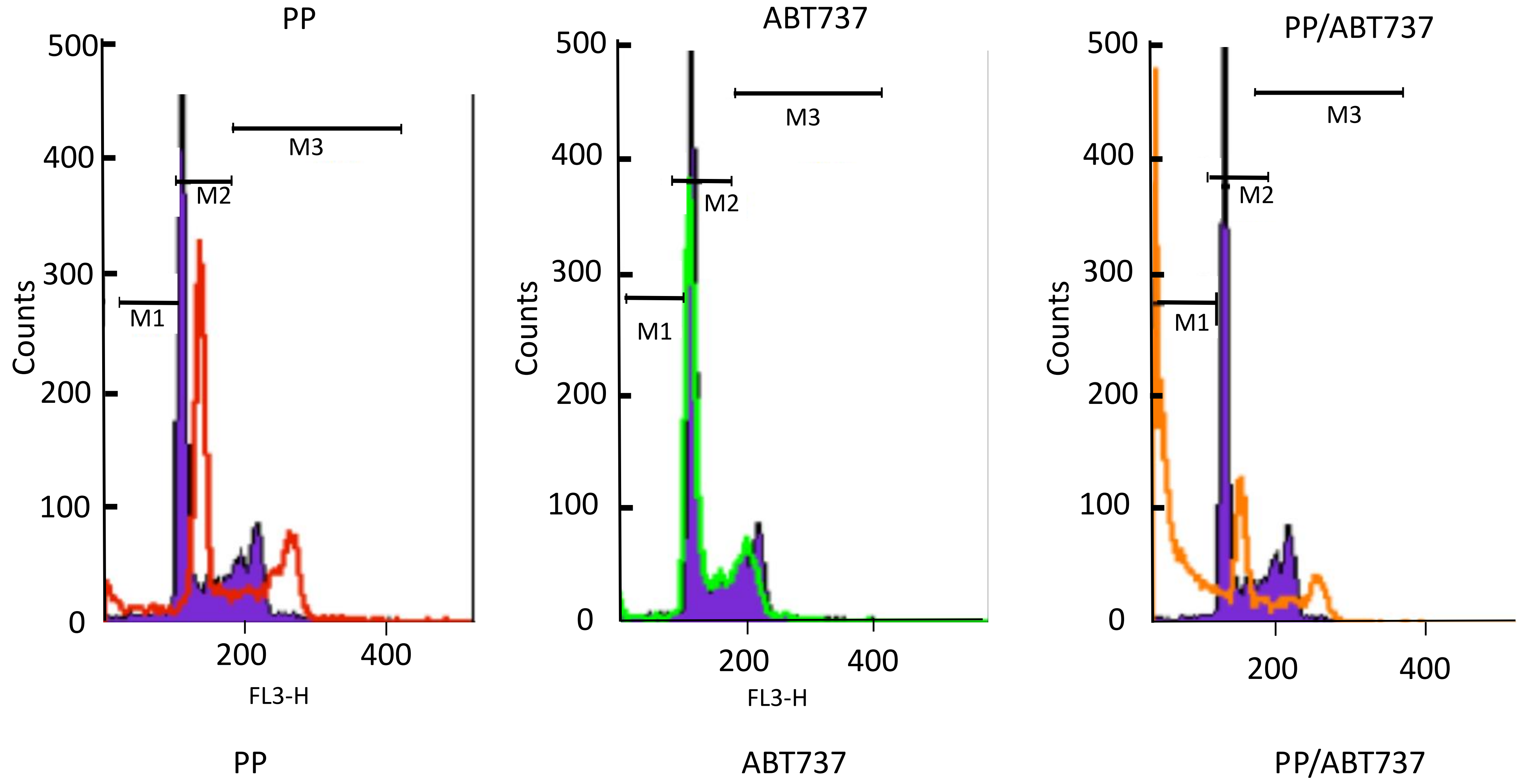


**Supplementary Figure 6** Treatment of SW620 cells with PP/ABT737 induces cell death via apoptosis

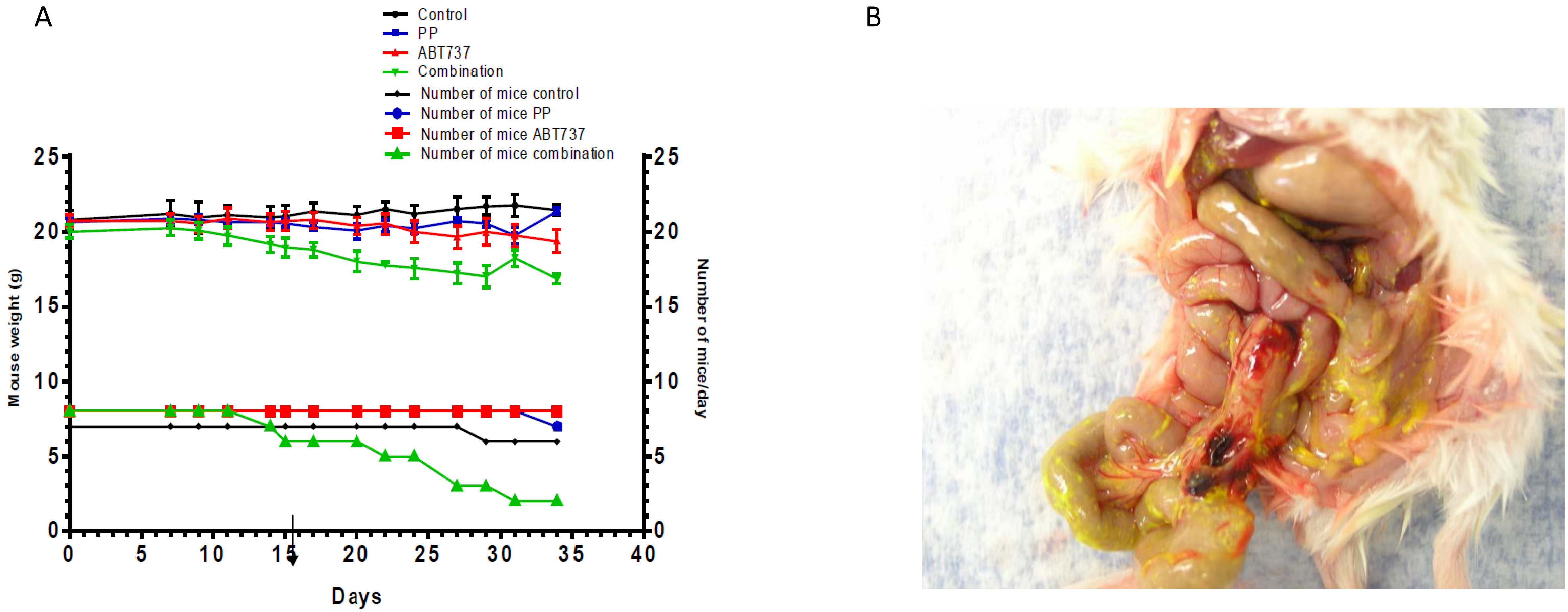


ight.

**Supplementary Figure 7**: A, Treatment of mice with the pyrvinium pamoate and ABT737 combination caused weight loss in mice; B, ABT737 injected intraperitoneally precipitated in the peritoneum (yellow precipitate).

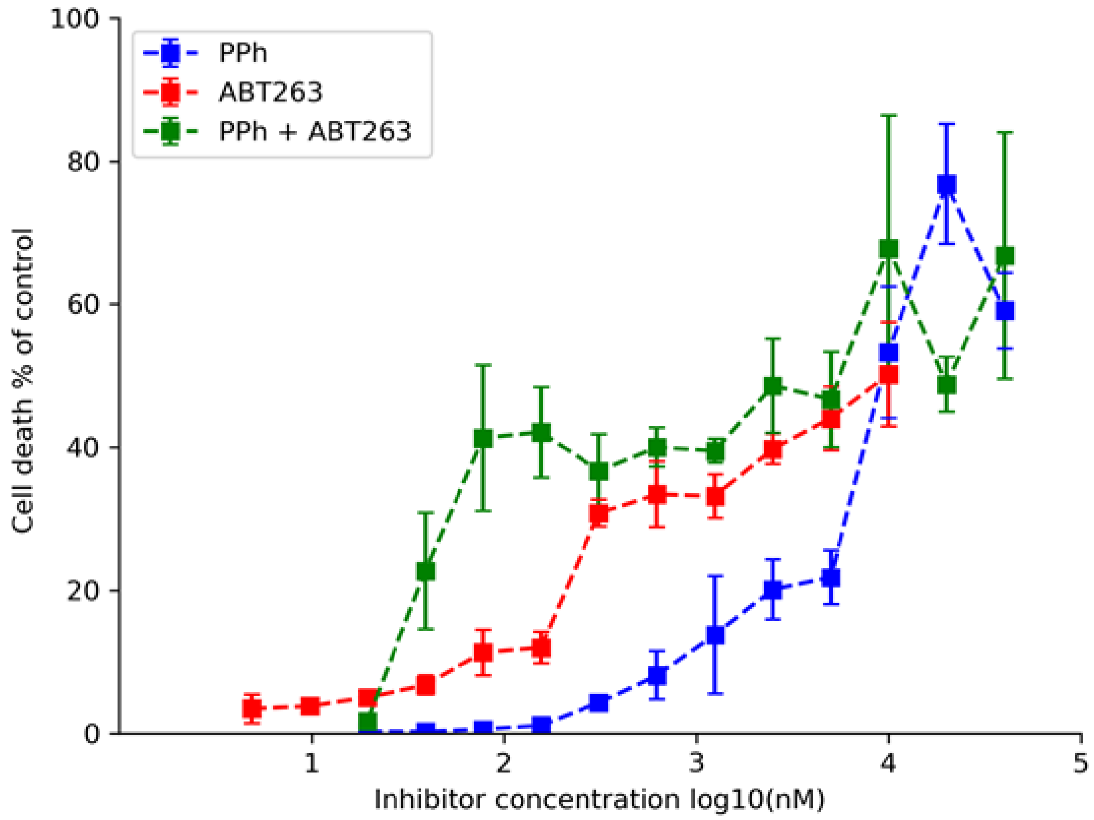


**Supplementary Figure 8**: Cytotoxicty effects of treatment with different concentrations of purvinium phosphate (PPh), ABT263 and the PPh/ABT263 (0.3um) combination on the CRC cell line LIM2405 growing in 3D-hanging drop cultures.

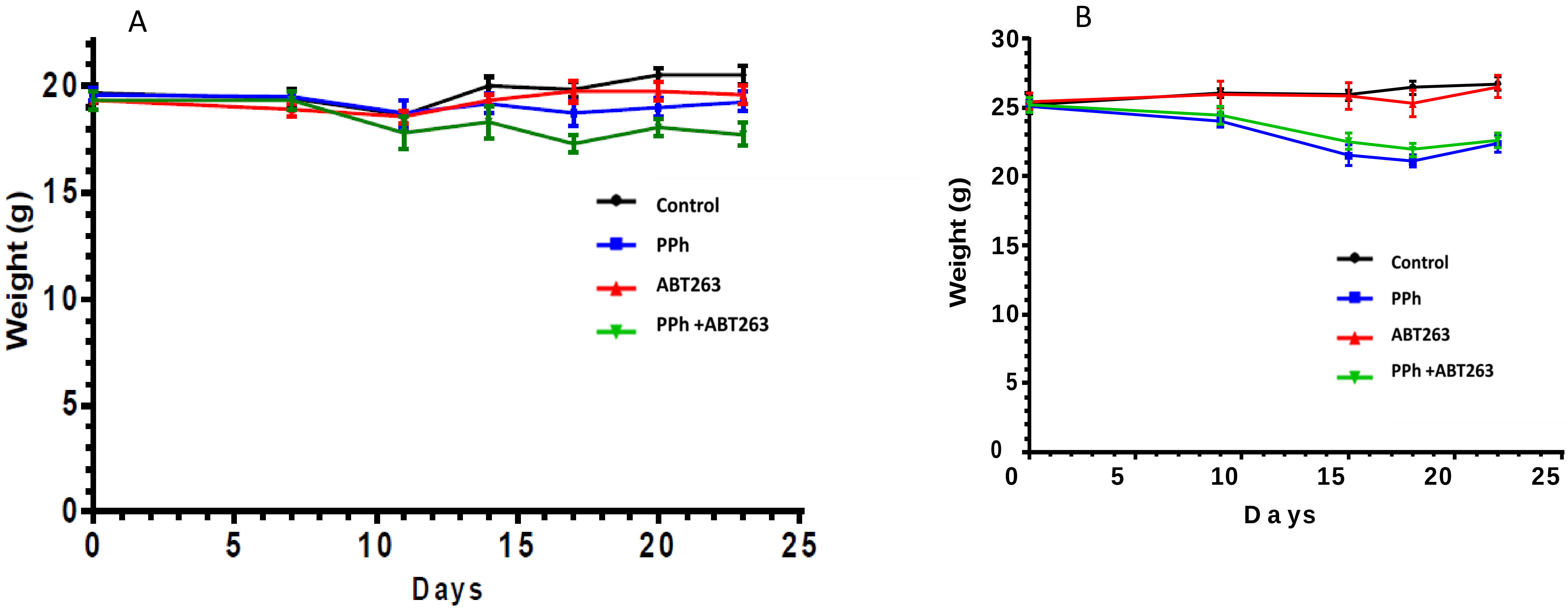


**Supplementary Figure 9** A, Effect on Body Weight of treating **female** mice carrying SW620 xenografts with Pyrvinium phosphate (PPh) and ABT263. The mice were weighed twice each week. The combination group appeared to lose weight from day 8 to 10. The food for all of the mice was then supplemented with Sustagen; B Effect on Body Weight of treating **male** mice carrying SW620 xenografts with Pyrvinium phosphate (PPh)and ABT263. The mice were weighed twice a week. Two groups (PP and combination) appeared to lose weight from day 15 The food for all of the mice was then supplemented with Sustagen.
